# Oxidized Lipids and CD36-Mediated Lipid Peroxidation in CD8 T Cells Suppress Anti-Tumor Immune Responses

**DOI:** 10.1101/2020.09.03.281691

**Authors:** Shihao Xu, Omkar Chaudhary, Patricia Rodríguez-Morales, Xiaoli Sun, Roberta Zappasodi, Ziyan Xu, Antonio F. M. Pinto, April Williams, Dan Chen, Jun Siong Low, Yagmur Farsakoglu, Wenxi Tang, Haiping Wang, Siva Karthik Varanasi, Bryan McDonald, Victoria Tripple, Michael Downes, Ronald M. Evans, Nada A. Abumrad, Taha Merghoub, Jedd D. Wolchok, Maxim N. Shokhirev, Ping-Chih Ho, Joseph L. Witztum, Brinda Emu, Guoliang Cui, Susan M. Kaech

## Abstract

T cell metabolic fitness plays a pivotal role in anti-tumor immunity and metabolic deregulation causes T cell dysfunction (i.e., ‘exhaustion’) in cancer. We identify that the scavenger receptor CD36 limits anti-tumor CD8^+^ T cell effector functions through lipid peroxidation. In murine tumors, oxidized phospholipids (OxPLs) were highly abundant and CD8^+^ TILs increased uptake and accumulation of lipids and lipid peroxidation. Functionally ‘exhausted’ CD8^+^ TILs substantially increased CD36 expression and CD36-deficient CD8^+^ TILs had more robust anti-tumor activity and cytokine production than wild-type cells. We further show that CD36 promotes uptake of oxidized low-density lipoproteins (OxLDL) and induces lipid peroxidation in CD8^+^ TILs, and OxLDL inhibits CD8^+^ T cell functions in a CD36-dependent manner. Moreover, glutathione peroxidase 4 (GPX4) over-expression lowers lipid peroxidation and restores functionalities in CD8^+^ TILs. These results define a key role for an oxidized lipid-CD36 axis in promoting intratumoral CD8^+^ T cell dysfunction.

## Introduction

Breakthroughs in cancer immunotherapy have revealed the power of harnessing immunity to fight cancer. However, only a small fraction of patients respond to current immunotherapies because the tumor microenvironment (TME) is highly immunosuppressive, highlighting the need to identify additional critical barriers that repress anti-tumor immune responses. Metabolic transformation is one cardinal feature of cancers, and the TME is metabolically distinct from normal tissue or circulation (Buck et al., 2017; Sullivan et al., 2019). An increasing body of evidence suggests that nutrient availability in the TME is a key influence on immune responses. For example, depletion of glucose and tryptophan, as well as accumulation of lactate, free fatty acids (FFAs) and cholesterol in the TME contribute to immunosuppression (Brand et al., 2016; Chang et al., 2015; Ho et al., 2015; Ma et al., 2019; Platten et al., 2012; Zhang et al., 2017).

Increased lipid accumulation is becoming a recognized hallmark of the TME that promotes immunosuppression (Al-Khami et al., 2017; Ma et al., 2019; Zhang et al., 2017). Intratumoral immune cells appear to metabolically adapt by enhancing lipid uptake or storage, which has been linked to their dysfunction. For example, lipid accumulation blunts antigen presentation in intratumoral dendritic cells (Cubillos-Ruiz et al., 2015; Herber et al., 2010; Veglia et al., 2017), and induces suppressive functions in neutrophils or myeloid-derived suppressor cells in tumors by promoting uptake of arachidonic acid, PGE2 synthesis, and fatty acid oxidation (FAO) (Al-Khami et al., 2017; Hossain et al., 2015; Veglia et al., 2019; Yan et al., 2019). Tumor-associated macrophages increase lipid accumulation and in an *in vitro* co-culture system, tumor cells induce neutral lipid storage and FAO in macrophages (Su et al., 2020). Additionally, accumulation of very long chain fatty acids or cholesterol has been shown to induce CD8^+^ TIL dysfunction (i.e., reduced cytotoxicity and TNF and IFNγ cytokine production), which is commonly referred to as T cell “exhaustion” (Ma et al., 2019; Manzo et al., 2020; Schietinger et al., 2016; Zajac et al., 1998). Lastly, increased lipid uptake in intratumoral regulatory CD4^+^ T cells (Tregs) promotes their maintenance and suppressive functions (Pacella et al., 2018; Wang et al., 2020). This in part was shown to involve the scavenger receptor CD36, also known as FAT, which is an important transporter of FFAs and oxidized lipids (such as oxidized LDL (OxLDL) and phosphocholine containing phospholipids (referred to herein as OxPLs)), and plays a major role in atherosclerosis (Abumrad et al., 1998; Boullier et al., 2005; Hajri and Abumrad, 2002; Que et al., 2018).

Deregulated lipid metabolism and increased reactive oxygen species (ROS) production can lead to lipid peroxidation, which has been implicated in a plethora of pathological settings including cardiovascular disease and cancer (Dixon and Stockwell, 2014; Yang and Stockwell, 2016). OxPLs and OxLDL underlie atherosclerosis, in part by promoting CD36-mediated uptake of OxLDL, inflammatory gene expression and generation of foamy macrophages (Que et al., 2018; Witztum and Steinberg, 1991). Increased lipid peroxidation in cancer cells has started to receive substantial attention because inactivation of cystine/glutamate antiporter xCT or glutathione peroxidase 4 (GPX4) can sensitize cancer cells to ferroptosis, a programmed cell death initiated by enhanced lipid peroxidation (Dixon et al., 2012; Dixon et al., 2014; Ingold et al., 2018; Yang et al., 2014). Interestingly, CD8^+^ T cells can regulate lipid peroxidation-dependent ferroptosis in tumor cells upon by secreting IFNg upon α-PD-L1 immunotherapy (Wang et al., 2019). GPX4-mediated regulation of lipid peroxidation also plays a vital role in maintaining periphery homeostasis and antigen-stimulated proliferation of T cells (Matsushita et al., 2015). However, the role of lipid peroxidation in antitumor immunity, particularly the effector functions of CD8^+^ T cells, remains elusive.

Here we show that OxPLs are a prevalent feature of the lipid-laden TME and CD8^+^ tumor infiltrating lymphocytes (TILs) up-regulate the expression of CD36, which increases their uptake of OxLDL and OxPLs and rate of lipid peroxidation. CD36 expression is enriched in the dysfunctional PD-1^+^ TIM-3^+^ TIL subset that displays reduced IFNg and TNF production. Genetic ablation of CD36 in CD8^+^ TILs increases their effector functions and results in enhanced tumor control. Importantly, CD36-deficient CD8^+^ TILs display reduced OxLDL uptake and lipid peroxidation. In addition, OxLDL inhibits CD8^+^ T cell cytokine production in a CD36-dependent manner. Suppressing lipid peroxidation via GPX4 over-expression can boost CD8^+^ TILs effector functions and enhance tumor control. Collectively, these findings uncover a new mode of immunosuppression in the TME by showing that elevated lipid oxidation— a common characteristic of tumor cells— also leads to increased oxidized lipid import by CD8^+^ TILs that in turn causes greater lipid peroxidation and T cell dysfunction. This study also discovers that CD36 plays a major role in this process, further underscoring the therapeutic potential of blocking CD36 to boost anti-tumor immunity.

## Results

### CD8 TILs adapt to increased lipid abundance in the TME

Several reports have shown that immune cells often have increased lipid content associated with abnormal functions in tumors, which could arise via increased de novo lipogenesis or import (Al-Khami et al., 2017; Ma et al., 2019; Manzo et al., 2020; Veglia et al., 2019; Wang et al., 2020; Zhang et al., 2017). To better understand what types of lipids immune cells are exposed to and could potentially import in tumors, we profiled the composition and abundance of various lipids within tumor interstitial fluid (TIF), which can be considered a proxy of the ‘extracellular milieu’ of the TME, from B16 or MC38 implanted tumors. When compared to serum from the same animal, the TIF contained greater amounts of many species of free fatty acids (FFAs), as shown previously (Ma et al., 2019; Zhang et al., 2017), as well as acyl-carnitines, ceramides, and esterified cholesterol (**Fig 1A**). Likely in response to increased lipid availability in tumors, we observed that CD8^+^ TILs bound to and/or imported more cholesterol and long- and medium-chain free fatty acids (FFAs) than CD8^+^ T cells in the spleen based on flow cytometric analysis of T cells cultured with fluorescently conjugated lipid substrates (NBD cholesterol, Bodipy C12, and Bodipy C16). Additionally, the CD8^+^ TILs exhibited overall higher neutral lipid content (e.g., TAGs and CEs based on Bodipy 493) compared to their splenic counterparts (**Fig 1B, Fig S1A**). Altogether, these results indicate that intratumoral CD8^+^ T cells adapt to the lipid-laden TME by enhancing uptake and storage of fatty acids and cholesterol.

**Figure 1.**
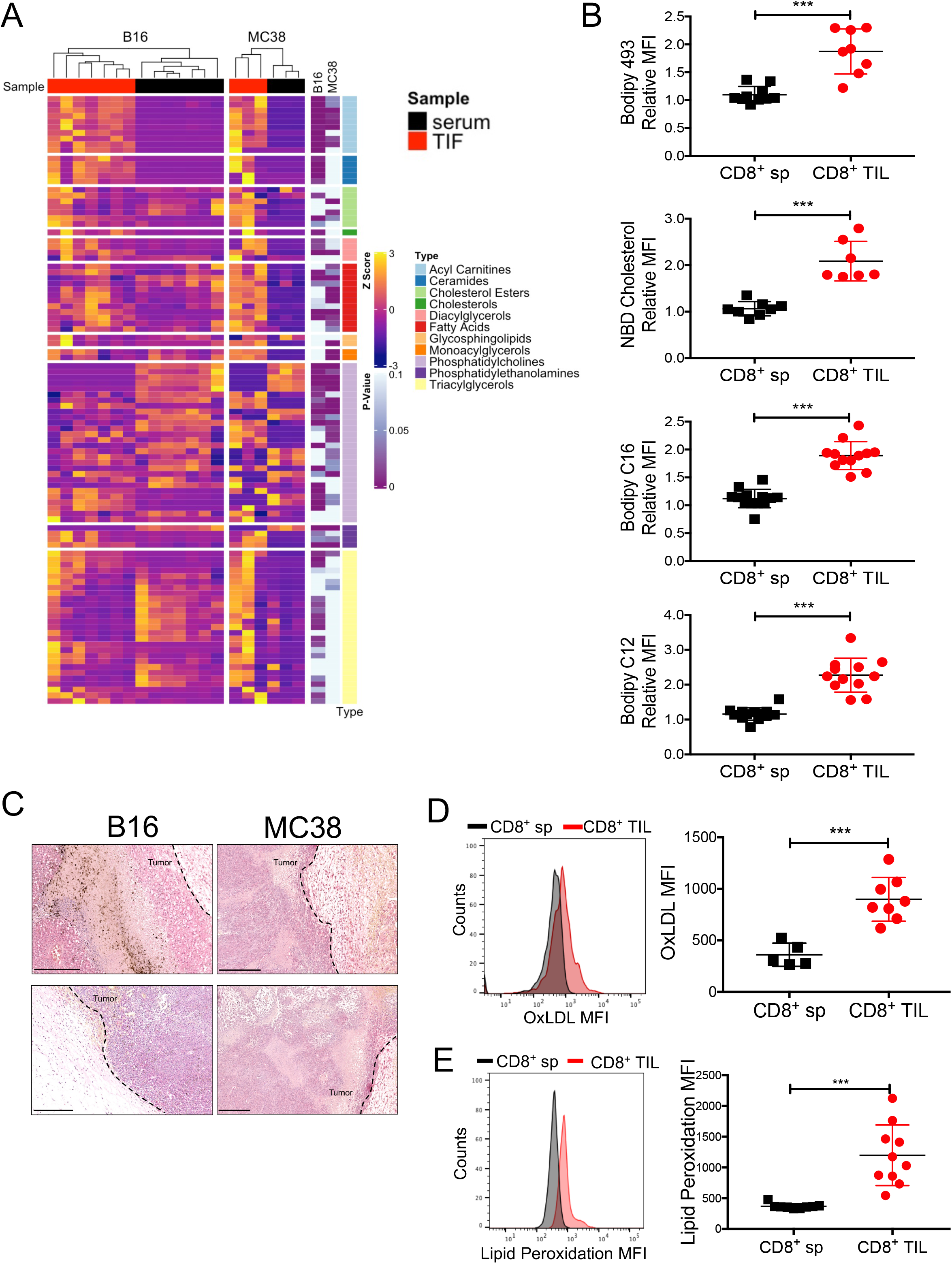
Increased exposure of CD8^+^ TILs to oxidized lipids in the TME. C57BL/6J mice were implanted with B16 or MC38 cells and tumors or splenocytes were examined 21 days later. (A) Lipids in the tumor interstitial fluids (TIF) from B16 or MC38 tumors were measured by mass spectrometry and compared to those in the sera. Heatmap shows the relative abundance of each lipid species normalized to protein concentration (shown as row Z-score). Results are representative of two experiments. (B) Neutral lipid content (Bodipy 493), uptake of cholesterol and fatty acids (NBD cholesterol, Bodipy C12, and Bodipy C16), were compared between CD8^+^ T cells isolated from the spleen (CD8^+^ sp) or B16 tumors (CD8^+^ TILs) using flow cytometry. Congenic Thy1.1 naïve splenocytes were spiked into each well to control for sample-to-sample variation and serve as an internal reference for normalizing lipid staining. Relative Mean Fluorescent Intensity (MFI) was calculated as the MFI ratio between the CD8^+^ T cells in the spleen or tumor relative to those in the internal reference. (C) Oxidized phospholipids (OxPLs) were stained in B16 and MC38 tumors with the biotinylated E06 antibody and detected with alkaline phosphatase conjugated avidin. The nuclei were stained with hematoxylin. Data are representative of three tumors/ group. Scale bar, 250 μm. (D) Uptake of oxidized LDL (OxLDL) in CD8^+^ sp or CD8^+^ TILs from B16 tumors was measured using fluorescently conjugated OxLDL and flow cytometry. (E) Lipid peroxidation in splenic CD8^+^ sp or CD8^+^ TILs from B16 tumors was quantified using BODIPY™ 581/591 C11 and flow cytometry. Data shown are mean ± SEM. Statistical analyses for (A, B, D, E) were performed by two-tailed unpaired Student’s t-test, ***p < 0.001. Samples were pooled from 2-3 experiments with each group containing n=7-12 (B), n=5-8 (D) or n=9-10 (E) animals.

Oxidative stress is a common feature of cancers (Reuter et al., 2010), and polyunsaturated fatty acids (PUFAs) are highly vulnerable to ROS-induced peroxidation (Yang et al., 2016). PUFA-derived OxPLs are not only found in OxLDL, but are prominent on apoptotic cells, apoptotic bodies and vesicles, and necrotic tissue (Binder et al., 2016), all of which are common in tumors. To look at the extent of OxPLs present in tumors, we performed immunohistochemistry (IHC) on B16 and MC38 tumor sections with the E06 natural antibody (NAb) that recognizes OxPLs (Shaw et al., 2000) (**Fig 1C, Fig S1B**). This revealed intense and diffuse OxPLs staining in both tumor types relative to adjacent healthy skin, demonstrating the prevalence of OxPLs in the TME. OxLDL is another abundant source of OxPL in tissues and we found that CD8^+^ TILs had higher rates of OxLDL uptake relative to splenocytes using fluorescently conjugated OxLDL and flow cytometry (**Fig 1D**). Importantly, we showed that CD8^+^ TILs also displayed a 2-4 fold increase in lipid peroxidation than their counterparts in the spleen based on the BODIPY^®^ 581/591 C11 lipid peroxidation assay (**Fig 1E**). This likely results from increased import of oxidized lipid substrates and/or enhanced intracellular lipid peroxidation due to imbalances in redox states and enhanced ROS in CD8^+^ TILs.

### CD36 is expressed on functionally exhausted TILs

The increase in lipid uptake and OxLDL binding by CD8^+^ TILs motivated us to look more closely at expression of specialized lipid transporters on the TILs, such as CD36. We found that mRNAs for *Cd36* and other FFA transporters including *Fabp4* and *Fabp5*, progressively increased as CD8^+^ TILs became more exhausted in murine liver cancer (GSE89307) (Philip et al., 2017) (**Fig S2A**). Thus, we hypothesized that functionally exhausted CD8^+^ TILs may up-regulate CD36 to metabolically adapt to heightened lipid availability in the TME. In support of this idea, we observed that CD8^+^ TILs from B16 or MC38 tumors had increased CD36 expression in ~20-40% of the cells (**Fig 2A**). Consistent with a recent discovery that CD36 maintains survival and suppressive functions of intratumoral Tregs (Wang et al., 2020), we observed that intratumoral CD4^+^ T cells, especially FOXP3^+^ Tregs, also increase CD36 expression (**Fig S2B**). In addition, the expression of CD36 on CD8^+^ TILs from early-stage tumors was barely detectable compared to those in late-stage tumors, suggesting CD36 expression is progressively acquired on CD8^+^ TILs as tumors grow (**Figs S2C-D**). Given that T cell exhaustion follows a differentiation path defined by progressive acquisition of inhibitory receptors (PD-1 and TIM-3) and granzyme B expression and loss of proinflammatory cytokine (TNF and IFNg) secretion (McLane et al., 2019; Wherry and Kurachi, 2015), we further separated CD8^+^ TILs based on PD-1 or TIM-3 expression into an effector cell subset (PD-1^-^ TIM-3^-^), an intermediate exhausted subset (PD-1^+^ TIM-3^-^), and a terminally exhausted subset (PD-1^+^ TIM-3^+^). Notably, TNF production negatively correlated with TIM-3 and TOX expression, and was progressively lost from the effector ➔ intermediate ➔ terminally exhausted subset (**Figs S2E-F**). CD36 was most highly expressed on the terminally exhausted subset (PD-1^+^ TIM-3^+^) of CD8^+^ TILs in B16 tumors or MC38 tumors (**Fig 2B**) and most of the CD36^+^ TILs expressed the immune-suppressive cytokine IL-10 based on the Thy1.1^+^ IL-10 reporter mice (Jin et al., 2010) (**Fig 2C**). These data demonstrated that CD36^+^ TILs were in a more functionally exhausted state. Importantly, the CD36^+^ PD-1^+^ CD8^+^ TILs displayed the highest amount of LDL, OxLDL, cholesterol, FFA uptake and neutral lipid content (Bodipy 493) relative to the PD-1^+^ CD36^-^ and PD-1^-^ CD36^-^ TILs (**Fig 2D**).

**Figure 2.**
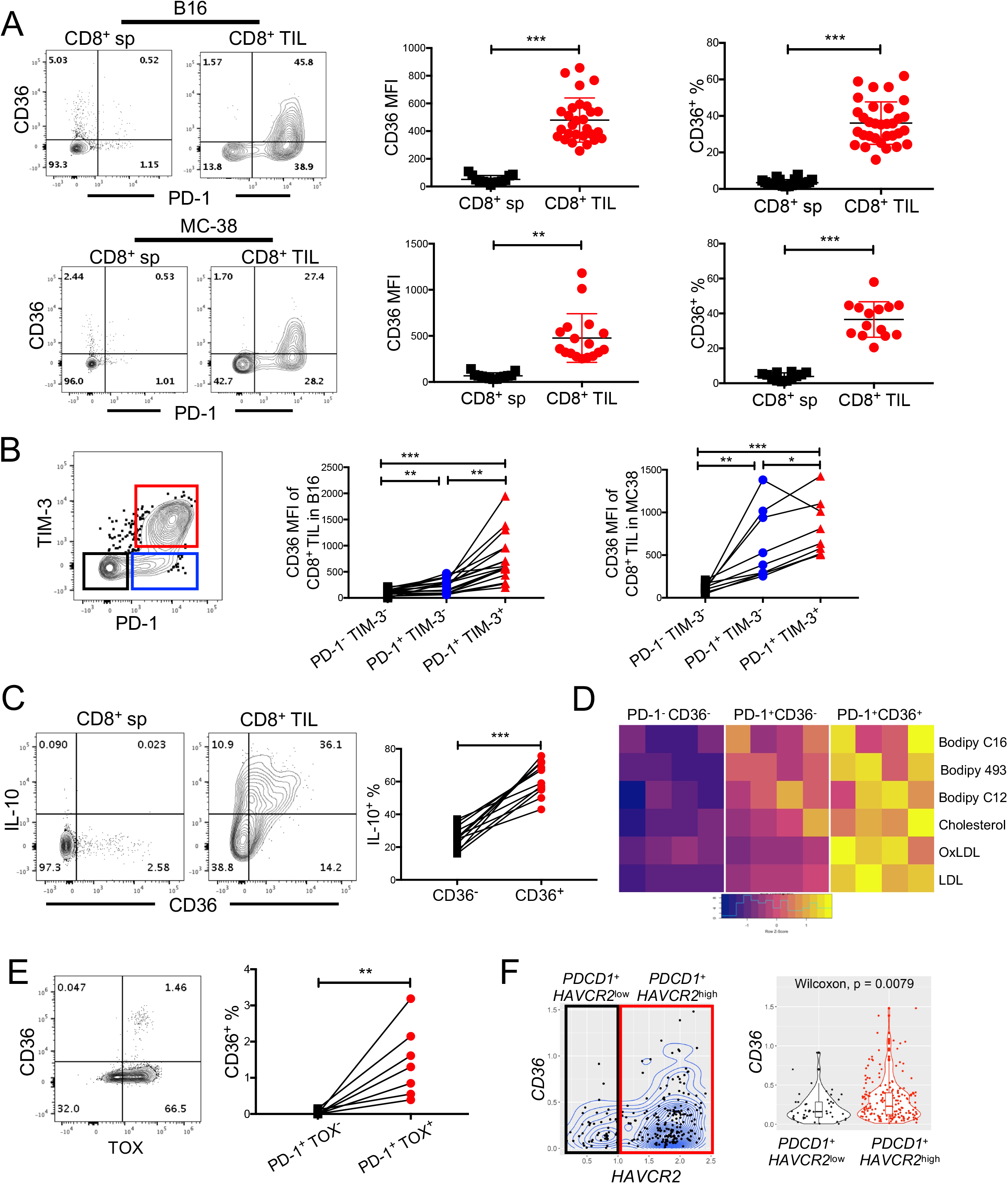
CD36 is expressed on functionally exhausted CD8^+^ TILs. (A, B, D) C57BL/6J mice were implanted with B16 or MC38 tumor cells as indicated below and tumors or splenocytes were examined 21 days later. (A-B) The expression of CD36, PD-1 and TIM-3 was measured in CD8^+^ sp and CD8^+^ TILs from B16 tumors or MC38 tumors using flow cytometry. Contour plots show representative staining patterns and scatter plots show the cumulative MFI or % CD36^+^ of total CD8^+^ T cells (A) or PD-1^-^ TIM-3^+^, PD-1^+^ TIM-3^-^ or PD-1^+^ TIM-3^+^ subsets of CD8^+^ TILs (B). (C) IL-10-reporter mice (10Bit) were implanted with B16 tumor cells and 21 days later CD36 and IL-10 expression (based on Thy1.1 staining) was assessed on CD8^+^ sp and CD8^+^ TILs using flow cytometry. Within each tumor analyzed, the percent of IL-10-expressing cells in the CD36^-^ or CD36^+^ CD8^+^ TIL subsets is shown. (D) As in Fig. 1, neutral lipid content (Bodipy 493) and uptake of polar lipids (Bodipy C12, Bodipy C16), cholesterol (NBD-Cholesterol), OxLDL or LDL were measured in the PD-1^-^ TIM-3^-^, PD-1^+^ TIM-3^-^ or PD-1^+^ TIM-3^+^ subsets of CD8^+^ TILs from B16 tumors using flow cytometry. Heatmap shows the MFI for each molecule analyzed (shown as row Z-score). (E) The expression of CD36 and TOX was measured by flow cytometry in PD-1^+^ CD8^+^ TILs from human melanomas. Within each tumor analyzed, the percent of CD36^+^ cells in the TOX^-^ and TOX^+^ subsets of PD-1^+^ CD8^+^ TILs is shown. (F) CD8^+^ TILs from human melanomas (GSE72056, (Tirosh et al., 2016)) were examined via scRNAseq analysis for *CD36* and TIM-3 (*HAVCR2*) mRNA expression. The analysis was restricted to the cells whose log-normalized mRNA expression of *CD3D, CD8A, PDCD1, CD36* and *HAVCR2* were all greater than 0. Contour plot (left) and boxplot (right) shows *CD36* mRNA abundance (log-normalized) in CD8^+^ TILs expressing either higher or lower amounts of *HAVCR2. P*-value was calculated by Wilcoxon test. Data shown are mean ± SEM and statistical analyses were performed by two-tailed unpaired Student’s t-test (A), and two-tailed paired Student’s t-test (B, C, E), * p < 0.05, ** p < 0.01, ***p < 0.001.. Samples were pooled from 2-5 experiments with each group containing n=13-29 (A), n=9-17 (B), n=8 (C), n=4 (D) animals or n=7 (E) patients.

To investigate if CD36 is associated with more exhausted TILs in human tumors, we measured the expression of CD36, TOX, and PD-1 in cryopreserved CD8^+^ TILs isolated from melanomas. These experiments showed that PD-1^+^ TOX^+^ CD8^+^ TILs had significantly higher CD36 expression relative to PD-1^+^ TOX^-^ counterparts, albeit the frequency of CD36^+^ cells was lower than in murine tumors (**Fig 2E**). This reduction in CD36 may be due to loss of signal after freezing and thawing of human T cells (data not shown). To further characterize human CD8^+^ TILs, we reanalyzed publicly available singlecell RNA sequencing (scRNAseq) data of human melanoma infiltrating CD8^+^ T cells (Tirosh et al., 2016). These analyses identified a subpopulation of CD8^+^ TILs that co-expressed *CD36, PDCD1* (PD-1) and *HAVCR2* (TIM-3) (**Fig 2F, left**). *PDCD1*^+^ *HAVCR2*^high^ of CD8^+^ TILs express higher levels of *CD36* than *PDCD1*^+^ *HAVCR2*^low^ counterparts (**Fig 2F, right**). Together, these data show that CD36 is a feature of both murine and human exhausted CD8^+^ TILs.

### CD36 promotes CD8^+^ TIL dysfunction in tumors

Checkpoint blockade targeting inhibitory receptors such as PD-1 and CTLA-4 unleash anti-tumor functions of CD8^+^ T cells and leads to long-lasting therapeutic effects in a fraction of cancer patients (Wei et al., 2018). As CD36 expression is associated with terminally exhausted CD8^+^ T cell states, we hypothesized that CD36 ablation on CD8^+^ TILs may rejuvenate anti-tumor functions of CD8^+^ T cells. To examine this hypothesis, we first implanted B16 or MC38 tumor cells subcutaneously into *Cd36^-/-^* germline knockout (KO) and *Cd36*^+/+^ control (WT) mice and monitored tumor growth and TIL functions. This showed that both types of tumors grew slower in *Cd36^-/-^* KO mice, and *Cd36^-/-^* KO CD8^+^ TILs produced more TNF^+^ or co-produced TNF^+^ IFNγ^+^ (i.e., were more polyfunctional) compared to *Cd36*^+/+^ WT controls (**Figs 3A-B, Figs S3A-B**). Granzyme B (GZMB) expression was also elevated in *Cd36^-/-^* KO CD8^+^ TILs relative to *Cd36*^+/+^ WT TILs in B16 tumors, while not significantly different in MC38 tumors. Importantly, depletion of T cells via αCD4 + αCD8 monoclonal antibodies largely restored B16 tumor growth in *Cd36^-/-^* KO mice, indicating that T cells play a crucial role in the enhanced tumor control of *Cd36^-/-^* mice (**Fig 3C**).

**Figure 3.**
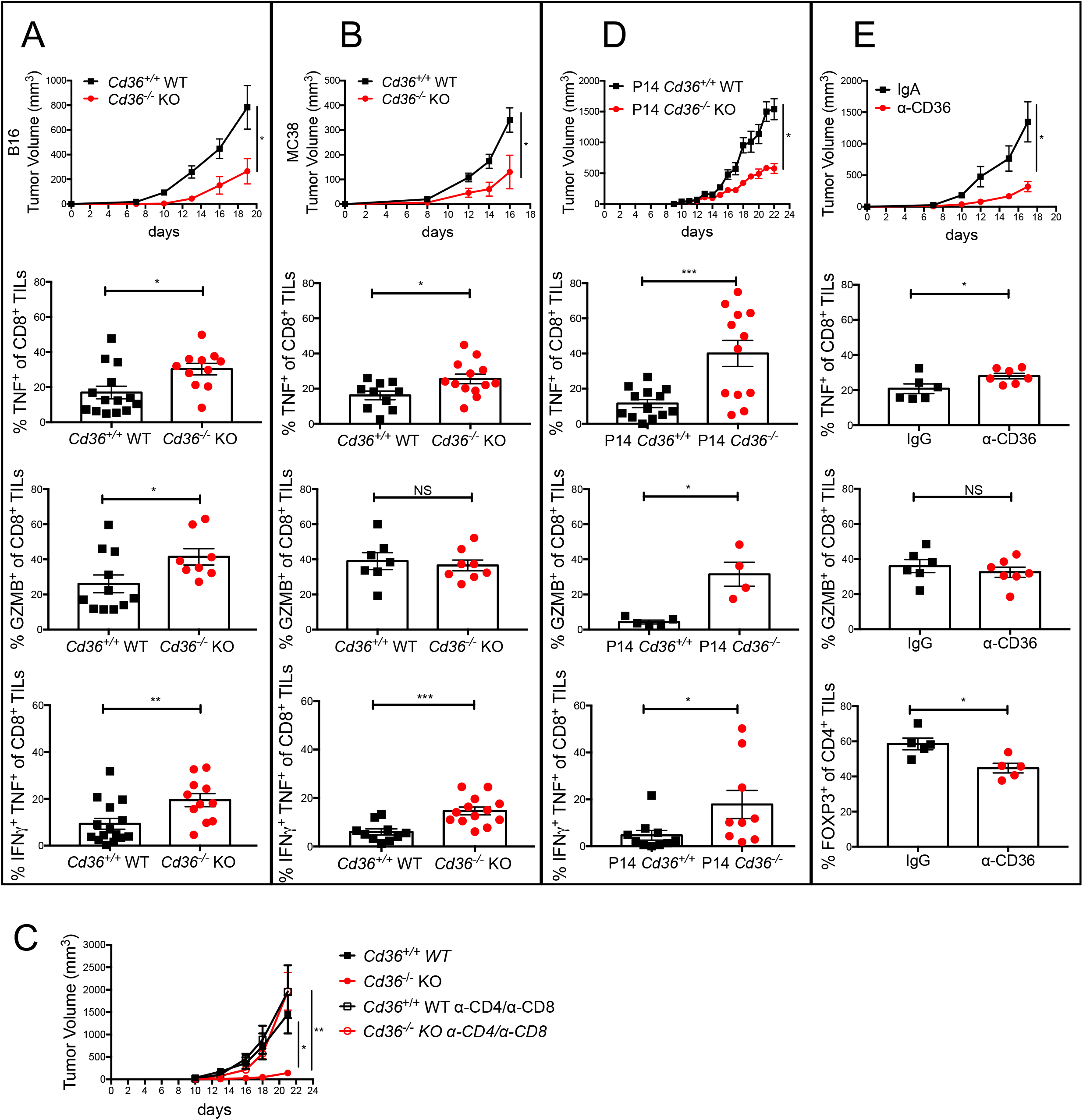
CD36 promotes CD8^+^ TIL dysfunction. (A-E) C57BL/6J mice were implanted with B16 or MC38 tumor cells as indicated below and tumors or splenocytes were examined 21 days later. (A-C) *Cd36*^+/+^ or *Cd36^-/-^* mice were implanted with B16 cells (A, C) or MC38 cells (B). Tumor growth was measured by caliper every 2-3 days and the expression of PD-1, TNF, IFNγ, and GZMB in CD8^+^ TILs was measured by flow cytometry. In (C), *Cd36*^+/+^ or *Cd36^-/-^* mice were treated with IgG or α-CD4/α-CD8 antibodies to delete T cells. (D) Mice were implanted with B16-gp33 cells and 10 days later 10^6^ P14 *Cd36*^+/+^ or *Cd36^-/-^* naïve CD8^+^ cells were adoptively transferred. Tumor growth was measured and the expression of TNF, IFNγ, and GZMB was measured in donor P14 CD8^+^ TILs. (E) Mice were implanted with B16 cells and treated with either IgA isotype control or α-CD36 Fab antibody (CRF D-2717, 200 μg, i.p., start on day 7 post tumor engraftment, every two days). Tumor growth was measured and the amounts of TNF and GZMB in CD8^+^ TILs and FOXP3 in CD4^+^ TILs were measured by flow cytometry. Data shown are mean± SEM, and statistical analyses were performed by two-tailed unpaired Student’s t-test, *p < 0.05; **p < 0.01; ***p < 0.001. Results were pooled from 2-4 experiments with each group containing n=7-12 (A), n=7-12 (B), n=6-7 (C), n=4-20 (D) or n=5-9 (E) animals.

Next, we examined the T cell-intrinsic function of CD36 in regulating anti-tumor functions by adoptively transferring naïve TCR transgenic P14 *Cd36*^+/+^ WT or *Cd36^-/-^* KO CD8^+^ T cells, which recognize the gp33-41 epitope from lymphocytic choriomeningitis virus (LCMV), into mice bearing B16-gp33 tumors that express the gp33 epitope. Similar to what was observed in the germline *Cd36^-/-^* KO mice, the B16-gp33 tumors grew slower in mice that contained P14 CD36^-/-^ KO T cells compared to P14 *Cd36*^+/+^ WT cells and P14 *Cd36^-/-^* KO TILs produced more TNF, IFNγ and GZMB than the *Cd36*-sufficient TILs (**Fig 3D, Fig S3C**). Consistently, our scRNAseq analysis of CD8^+^ TILs revealed that *Cd36^-/-^* KO cells relative to *Cd36*^+/+^ WT controls showed lower amounts of *S100a4* and *Nr4a2* mRNA, whose expression is elevated in exhausted CD8^+^ T cells (Chen et al., 2019; Haining et al., 2008), and higher expression of *Gzmb* and *Ifng*, in line with elevated effector functions. In addition, interferon induced genes (*Ifit3, Ifit1, Tnfsf4, Ifitm3, Isg15*) tended to be increased as well in *Cd36^-/-^* KO cells, but the difference for some of those genes was not significant at an FDR < 0.1. Altogether, this demonstrated that increased expression of CD36 on PD-1^+^ TIM-3^+^ CD8^+^ TILs suppressed their antitumor effector functions.

Lastly, we explored whether antibody-mediated blockade of CD36 (α-CD36) could increase anti-tumor functions of CD8^+^ TILs and decrease tumor burden by treating mice with an α-CD36 blocking antibody every three days from day 7 post tumor implantation (200 μg, intraperitoneal (i.p.)). This revealed that α-CD36 blockade suppressed B16 tumor growth, promoted TNF expression in CD8 TILs, and reduced the frequency of intratumoral Tregs (**Fig 3E**). This is consistent with a prior study showing blocking CD36 suppressed tumor growth and synergized with anti-PD-1 checkpoint blockade (Wang et al., 2020). Combined together, these data reveal that CD36 expression promotes CD8^+^ TIL dysfunction and can be a therapeutic target for cancer in multiple tumor models.

### CD36 mediates OxLDL uptake in CD8^+^ TILs

In discovering that CD36^+^ PD-1^+^ CD8^+^ TILs displayed the highest amount of LDL and OxLDL uptake and accumulation of lipids (**Fig 2D**), we next asked if these properties were dependent on CD36. To this end, we compared the neutral lipid content and uptake of labeled cholesterol or FFAs between *Cd36*^+/+^ WT and *Cd36^-/-^* KO CD8^+^ TILs, but did not observe any significant differences, albeit there was a trend for lower neutral lipid content (**Figs S4A-D**). Nor did we find any differences in mitochondria potential or rates of fatty acid oxidation (FAO) between *Cd36*^+/+^ WT and *Cd36^-/-^* KO CD8^+^ TILs (**Figs S4E-F**). These data suggest that the FAA transport function of CD36 was likely not involved in regulating TIL function. However, consistent with the known ability of CD36 to mediate uptake of OxLDL, but not native LDL, we observed a selective loss of OxLDL, but not LDL, uptake in *Cd36^-/-^* KO CD8^+^ TILs compared to the WT controls (**Figs 4A-B**). Imaging flow cytometry confirmed the colocalization of OxLDL and CD36 in CD8^+^ TILs (**Fig 4C** and **Fig S4G-H**). Together, these data indicate that CD36 is required for OxLDL uptake in CD8^+^ TILs and correlates with greater TIL dysfunction.

**Figure 4.**
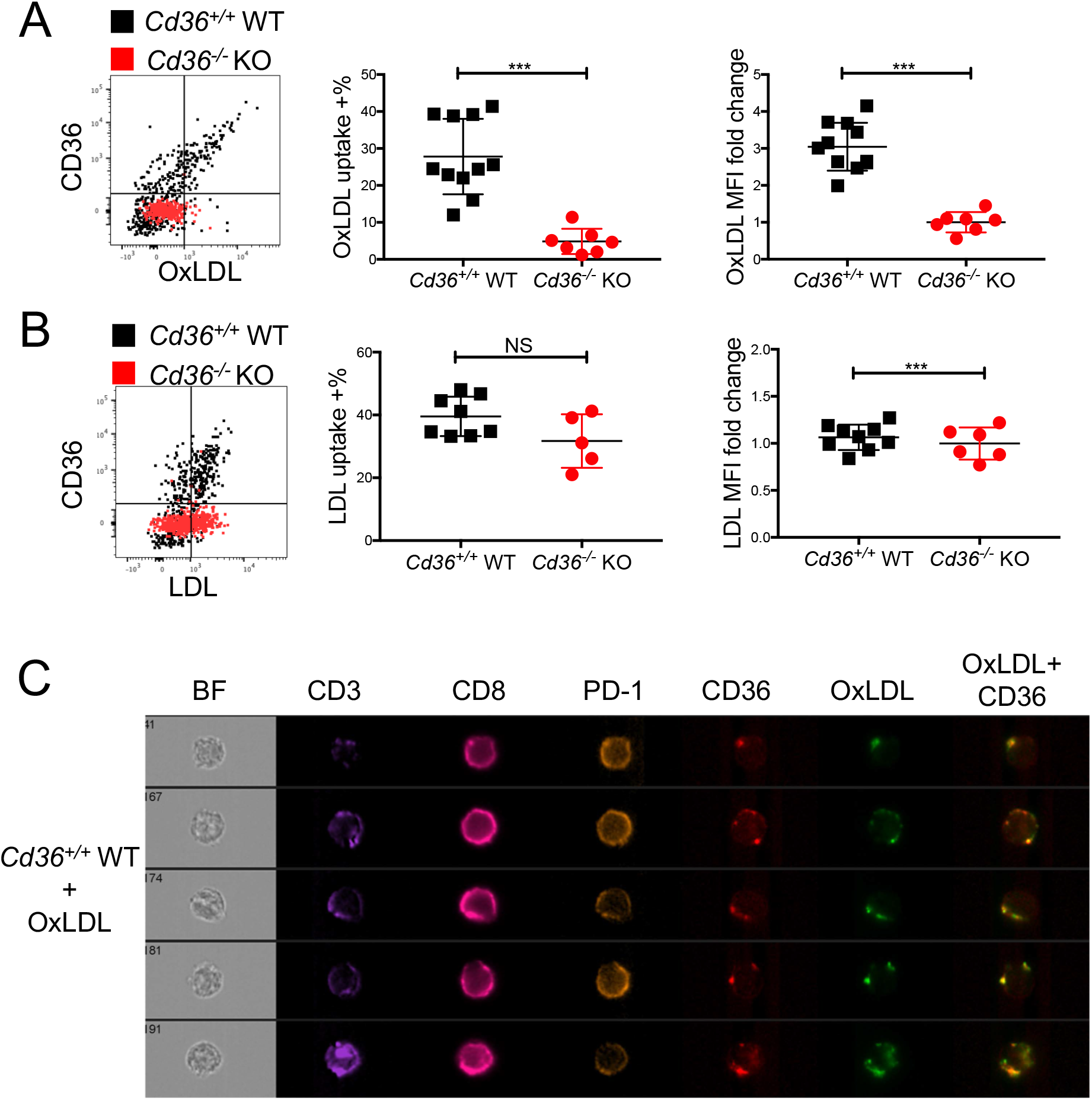
CD36 mediates OxLDL uptake in CD8^+^ TILs. (A-B) *Cd36*^+/+^ WT or *Cd36^-/-^* KO mice were implanted with B16 cells and the CD8^+^ TILs were isolated and stained with fluorescently conjugated anti-CD36 mAbs and OxLDL (A) or LDL (B) and analyzed by flow cytometry 21 days post tumor implantation. Note, the direct correlation between CD36 expression and OxLDL uptake on *Cd36*^+/+^ WT cells and lack of OxLDL, but not LDL, uptake in *Cd36^-/-^* TILs. (C) C57BL/6J mice were implanted with B16 tumor cells 21 days post implantation the expression of CD3, CD8, PD-1, CD36, and OxLDL uptake in *Cd36*^+/+^ CD8^+^ TILs were measured by Amnis ImageSteam^®^ flow cytometry. BF, bright field. Representative images are shown of 2 experiments. Note, the colocalization of CD36 and OxLDL in merged image (right most panel). Data shown are mean± SEM, and statistical analyses were performed by two-tailed unpaired Student’s t-test. ***p < 0.001, NS, non-significant. Results are pooled from 2-3 experiments with each group containing n=5-11 animals.

### OxLDL inhibits CD8 T cell effector function through CD36

Given that OxPLs were abundant in the TME and OxLDL is greatly enriched in OxPLs, we next wanted to examine the effects of increased oxidized lipid import on CD8^+^ TIL functions. We treated murine splenic CD8^+^ T cells activated *in vitro* with OxLDL, LDL, HDL, or sulfosuccinimidyl oleate (SSO), an irreversible inhibitor of CD36 (Kuda et al., 2013). This showed that OxLDL, but not LDL or HDL, suppressed TNF and IFNg production in CD8^+^ T cells and interestingly, SSO interfered with this inhibitory effect of OxLDL on effector functions *in vitro* (**Fig 5A**). Similarly, we found that OxLDL profoundly inhibited secretion of TNF and IFNg in human CD8^+^ T cells in vitro, which can be fully rescued by SSO (**Fig 5B**). Note, OxLDL did not substantially decrease cell viability in these culture conditions (**Fig 5A and Figs S5A-B**). Moreover, OxLDL potently repressed cytokine production by CD8^+^ TILs *ex vivo*, and this was largely ameliorated in *Cd36^-/-^* CD8^+^ TILs (**Fig 5C**). Together, these data show that CD36-dependent OxLDL uptake inhibits effector functions of CD8^+^ T cells, suggesting that an oxidized lipid-CD36 axis serves as a metabolic checkpoint of effector CD8^+^ T cells in the TME.

**Figure 5.**
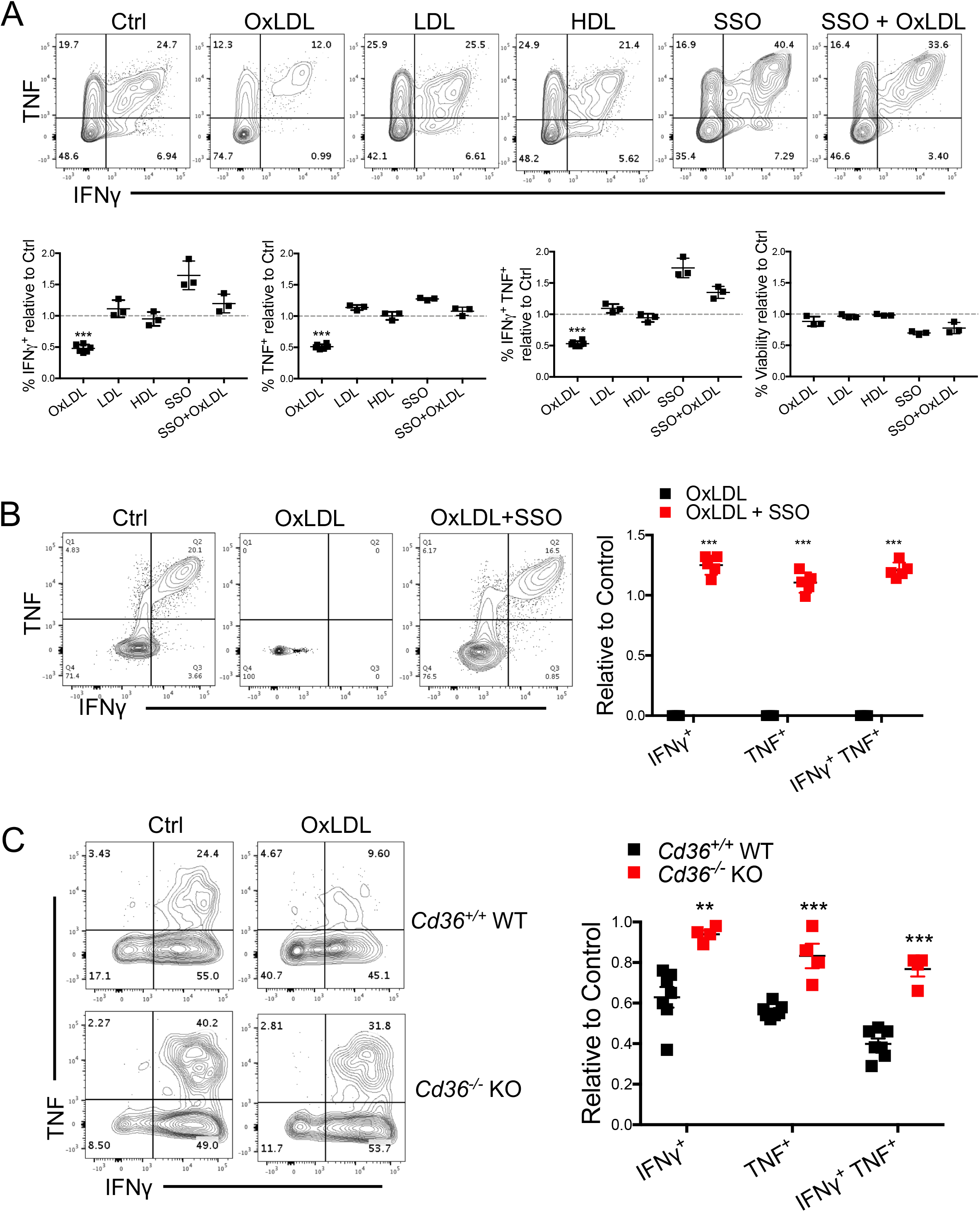
OxLDL inhibits CD8^+^ T cell function in a CD36-dependent manner. (A) P14 CD8^+^ T cells were activated *in vitro* with gp33 peptide plus IL-2 for 24 hrs and then treated with either vehicle control (Ctrl), OxLDL (50 μg/ml), LDL (50 μg/ml), HDL (50 μg/ml), SSO (100 μM), or the combination of OxLDL (50 μg/ml) and SSO (100 μM), for another 48 hrs. TNF, IFNγ and cell viability were then measured upon re-stimulation with gp33 for 6 hours and analyzed by flow cytometry. (B) Human PBMCs were treated with either vehicle control (Ctrl), OxLDL (50 μg/ml), or the combination of OxLDL (50 μg/ml) and SSO (100 μM). TNF, IFNγ and cell viability was measured by flow cytometry 16 hrs after stimulation with Staphylococcal enterotoxin B (SEB). (C) *Cd36*^+/+^ or *Cd36^-/-^* CD8^+^ TILs isolated from B16 tumors 21 days post implantation were purified by FACS and treated with either vehicle control (Ctrl) or oxLDL (50 μg/ml) for 24 hrs. TNF and IFNγ was measured by flow cytometry 4 hrs after stimulation with PMA/ionomycin. Data shown are mean± SEM and statistical tests were performed by two-tailed unpaired Student’s t-test (A, C), and two-tailed paired Student’s t-test (B), **p < 0.01, ***p < 0.001. Results are pooled from 23 experiments with each group containing n=3-6 animals (A, C) or n=5 healthy controls (B).

### OxLDL induces lipid peroxidation in CD8^+^ T cells through CD36

OxLDL and OxPLs induce oxidative stress in macrophages and endothelial cells, and promote lipid peroxidation in non-alcoholic fatty liver disease (Navab et al., 2004; Que et al., 2018; Sun et al., 2020; Witztum and Steinberg, 1991). Because we found that CD8^+^ TILs take up OxLDL in a CD36-dependent manner (**Fig 4B**), we hypothesized that this would increase lipid peroxidation. First, we measured the impact of OxLDL on lipid peroxidation of mouse or human CD8^+^ T cells activated *in vitro* using the BODIPY^®^ 581/591 C11 reagent and this revealed that OxLDL, but not LDL, enhanced lipid peroxidation in CD8^+^ T cells *in vitro* (**Figs 6A-B**). This suggested that OxLDL is sufficient to enhance lipid peroxidation in T cells, and in agreement with the increased OxLDL uptake and amounts of lipid peroxidation observed in CD8^+^ TILs relative to splenic CD8^+^ T cells (**Figs 1D-E**). Next, to examine whether CD36 could affect CD8^+^ TIL lipid peroxidation *in vivo* we compared the extent of lipid peroxidation between *Cd36*^+/+^ WT and *Cd36^-/-^* KO CD8^+^ TILs purified from murine B16 tumors, using the BODIPY^®^ 581/591 C11 reagent, and found CD36-deficient TILs displayed a lower level of lipid peroxidation (**Fig 6C, left**). Similarly, reduced lipid peroxidation was seen in P14 *Cd36^-/-^* KO TILs compared to *Cd36*^+/+^ WT cells, indicating that lipid peroxidation depends directly on T cell-intrinsic CD36 expression (**Fig 6C, right**). Collectively, these data indicate that CD36 is a major mediator of lipid peroxidation in CD8^+^ TILs.

**Figure 6.**
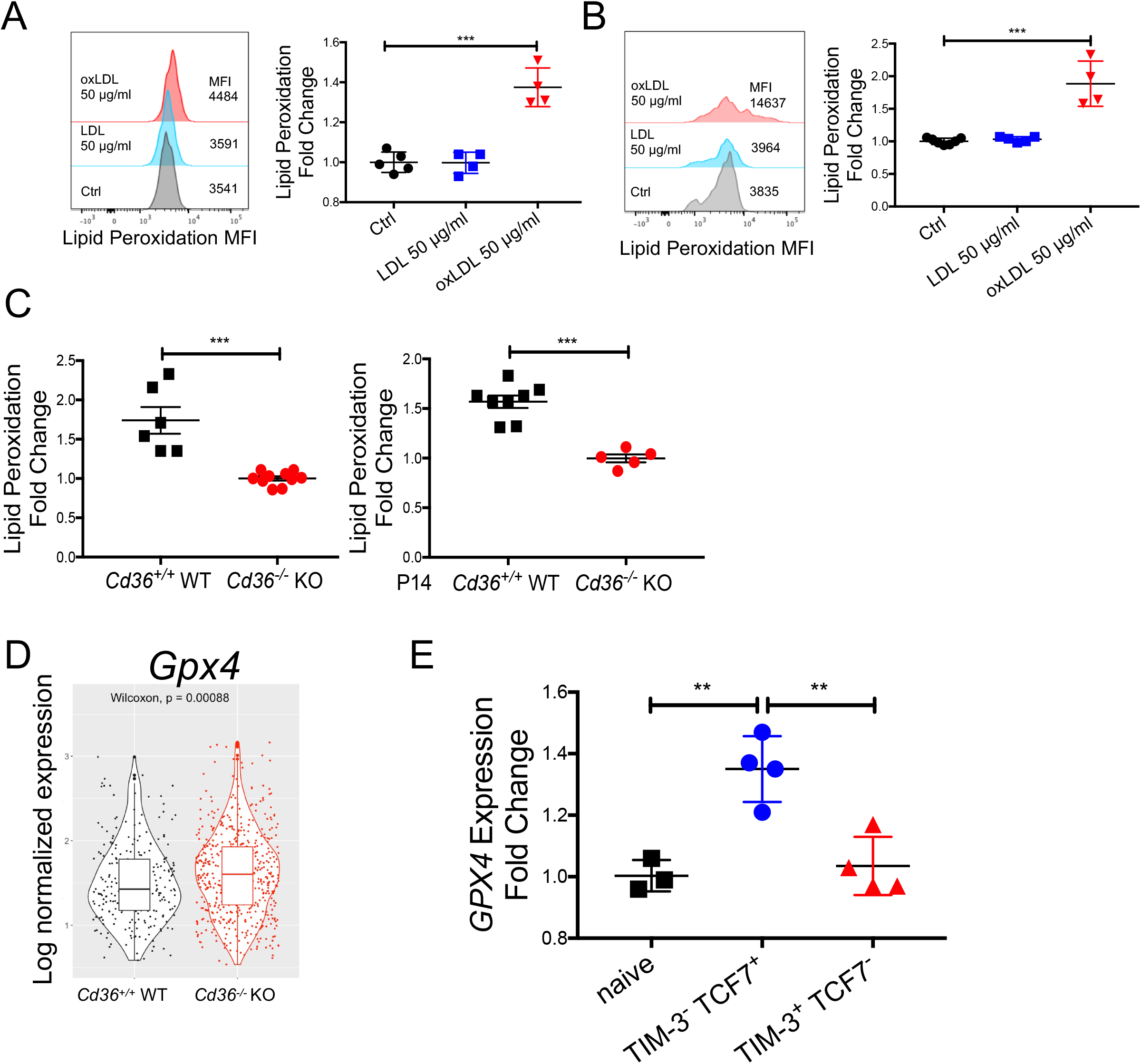
OxLDL induces lipid peroxidation in CD8^+^ T cells in a CD36-dependent manner. (A) P14 CD8^+^ T cells were activated *in vitro* for 24 hrs and then treated with vehicle control (Ctrl), OxLDL (50 μg/ml), LDL (50 μg/ml) for 24 hrs. The cells were then washed in PBS and incubated with BODIPY^®^ 581/591 C11 for lipid peroxidation assay. (B) Human PBMCs treated with vehicle control (Ctrl), LDL (50 μg/ml), oxLDL (50 μg/ml) for 24 hrs and then washed in PBS and incubated with BODIPY^®^ 581/591 C11 for lipid peroxidation assay. (C) BODIPY 581/591 C11 lipid peroxidation assay was performed directly *ex vivo* on CD8^+^ TILs isolated from B16 tumors implanted 21 days previously into *Cd36*^+/+^ WT or *Cd36^-/-^* KO mice (left) or B6 mice that contained P14 *Cd36*^+/+^ WT or *Cd36^-/-^* KO CD8^+^ TILs (right). Graphs show the fold change in BODIPY 581/591 C11 fluorescence in WT relative to KO TILs. (D) mRNA expression of *Gpx4* was compared between P14 *Cd36*^+/+^ WT or *Cd36^-/-^* KO TILs isolated from B16 tumors 21 days post implantation and analyzed by scRNAseq. *p*-value was calculated by Wilcoxon test. (E) Analysis of GSE114631 *Gpx4* mRNA expression in naïve, TIM-3^+^ TCF7^-^, or TIM-3^-^ TCF7^+^ P14 CD8^+^ TILs isolated from B16-gp33 tumors 6 days after tumors became palpable (Siddiqui et al., 2019). Data shown in A, B, C and E are mean ± SEM and statistical analyses were performed by two-tailed unpaired Student’s t-test, **p < 0.01, ***p < 0.001. Samples were pooled from 2-3 experiments with each group containing n=4-5 (A, B) and n=5-10 (C) animals.

### GPX4 over-expression restores CD8 T cell function in tumors

Lipid peroxidation can lead to ferroptosis, a unique form of programmed cell death that is characterized by iron overloading and increased lipid peroxides (Y ang and Stockwell, 2016). Glutathione peroxidase 4 (GPX4), a target of ferroptosis-inducing compounds (e.g., RSL3), can rescue cells from ferroptosis by degrading lipid peroxides (Ingold et al., 2018; Yang et al., 2014). GPX4 is not required for T cell development but is indispensable for T cell homeostasis and T-cell dependent immune responses (Matsushita et al., 2015). Correlating with reduced lipid peroxidation in *Cd36^-/-^* KO TILs, we observed a modest increase in *Gpx4* mRNA in these cells relative to the *Cd36*^+/+^ WT CD8^+^ TILs (**Fig 6D**). Further, *Gpx4* mRNA expression declined in CD8^+^ TILs from B16 tumors (GSE114631) as they differentiated from TCF7^+^ TIM-3^-^ progenitor-like to TCF7^-^ TIM-3^+^ terminally exhausted T cells (Siddiqui et al., 2019), suggesting that reduced *Gpx4* expression is associated with T cell exhaustion (**Fig 6E**).

Because CD36-dependent uptake of OxLDL induces lipid peroxidation and suppresses T cell functions, we reasoned that prevention of lipid peroxidation via GPX4 over-expression (OE), may rescue effector functions of T cells. To test this idea, we over-expressed GPX4 in P14 CD8^+^ T cells and performed adoptive T cell transfer to mice bearing B16-gp33 tumors. GPX4 OE in CD8^+^ TILs resulted in enhanced tumor control compared to the control cells (**Fig 7A**). Specifically, GPX4 OE led to marked increase in the number of CD8^+^ TILs (**Fig 7B**), and decreased lipid peroxidation in CD8^+^ TILs (**Fig 7C**). Moreover, GPX4 OE boosted anti-tumor effector functions of CD8^+^ TILs by increasing the secretion of TNF and IFNγ^+^ (**Fig 7D**). Together, our data suggest that suppression of lipid peroxidation could rescue anti-tumor functions of TILs, and GPX4 can be a novel target for CD8^+^ T cell-based immunotherapies.

**Figure 7.**
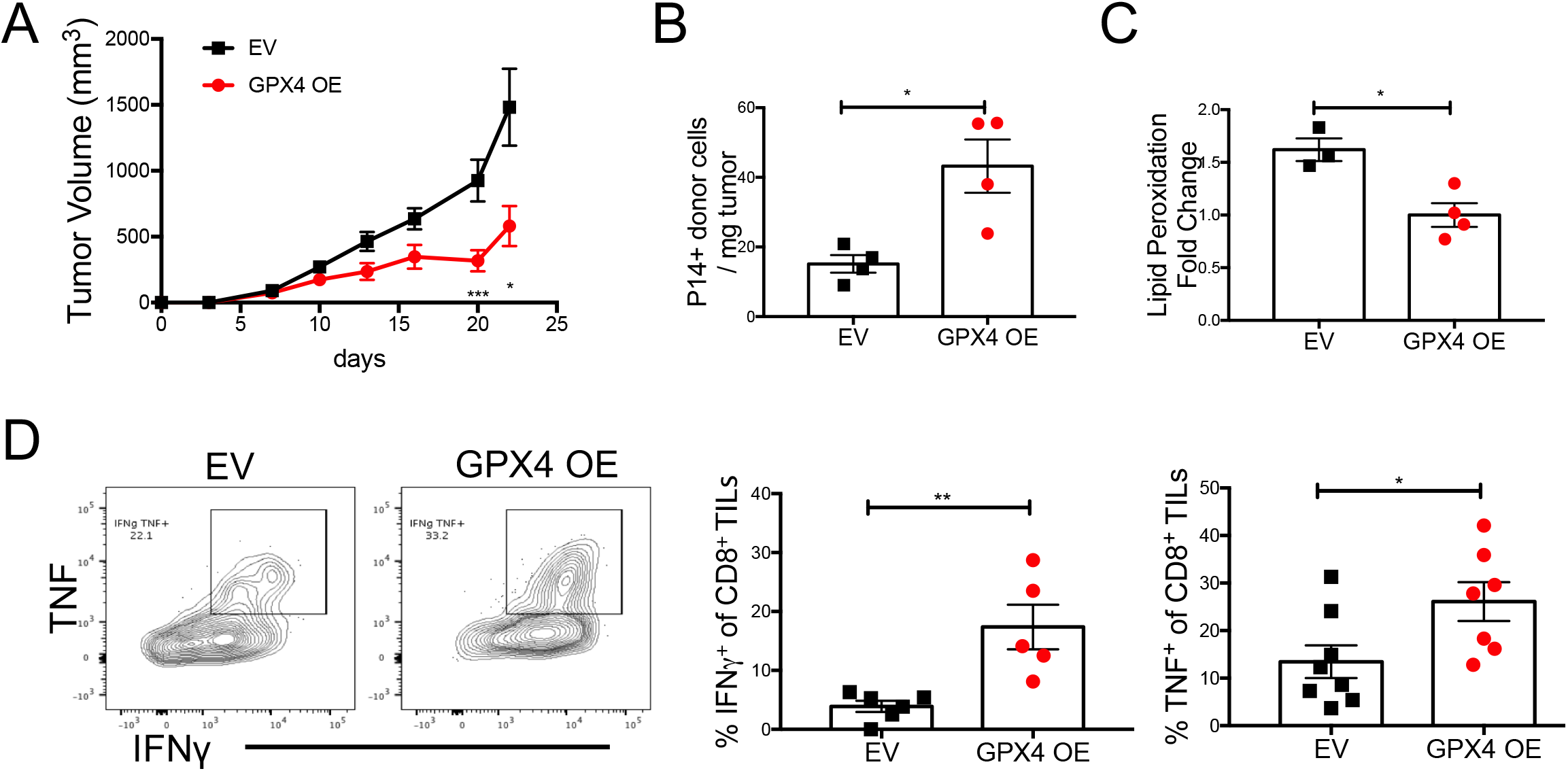
GPX4 OE restores CD8^+^ T cell function in tumors. (A-D) *In vitro* activated P14 CD8^+^ T cells were transduced with either empty retrovirus (EV) or a retrovirus overexpressing *Gpx4* (GPX4 OE) and 5×10^5^ cells were adoptively transferred into C57BL/6J mice previously implanted with B16 cells 7 days prior. Tumor growth was measured every 2-3 days and EV and GPX4 OE P14^+^ donor TILs were analyzed at day 21 post tumor implantation by flow cytometry for cell numbers (B), rate of lipid peroxidation (C) or IFNγ and TNF cytokine production (D). Data shown are mean± SEM and statistical analysis was performed by two-tailed unpaired Student’s t-test *p < 0.05, **p < 0.01, ***p < 0.001. Samples were pooled from 2 experiments with each group containing n=8 (A), n=3-4 (B, C), and n=5-7 (D) animals.

## Discussion

Identifying factors that cause immune suppression in the TME can lead to development of novel immunotherapies. While oxidized lipids are a common feature of inflamed tissues (Binder et al., 2016; Miller and Shyy, 2017), the role of oxidized lipids in the TME had not been well addressed before. Here our study suggests a new mode of immunosuppression in the TME: increased import of oxidized lipids by CD8^+^ TILs, likely caused by elevated lipid oxidation in tumors, leads to greater lipid peroxidation and dysfunction in CD8^+^ TILs. We found that ablation of CD36 or over-expression of GPX4, suppressed lipid peroxidation, boosted CD8^+^ TILs effector functions and enhanced tumor control. Our study illuminates immune modulatory effects of oxidized lipids in cancer.

Deregulated lipid metabolism is a common feature of the TME, and increased lipid uptake and accumulation is observed in many types of intratumoral immune cells, often associated with impaired anti-tumor immune function (Herber et al., 2010; Manzo et al., 2020; Su et al., 2020; Veglia et al., 2019; Wang et al., 2020; Zhang et al., 2017). Consistent with our findings, intratumoral CD8^+^ TILs in pancreatic cancers increase lipid deposition, specifically very-long chain fatty acids (VLCFAs) (Manzo et al., 2020). The metabolic fitness of intratumoral Tregs depends on enhanced CD36 expression for heightened lipid uptake and accumulation, and CD36-deficient Tregs had lower amounts of mitochondria specifically in tumors (Wang et al., 2020). In addition, *in vitro* co-culture with tumor cells enables macrophages to enhance neutral lipid storage and fatty acid oxidation in a CD36-dependent manner (Su et al., 2020). However, we did not observe significant differences in FFA analog uptake, neutral lipid accumulation, mitochondrial potential or fatty acid oxidation between *Cd36*^+/+^ and *Cd36*^-/-^ CD8^+^ TILs in B16 or MC38 tumors, likely highlighting cell type-specific roles for CD36. Given the redundant roles of different lipid transporters in lipid uptake and the importance of FABP4/5 for lipid homeostasis in the skin resident memory T cells (Pan et al., 2017), we postulate that FABP4/5 (and others) may be involved in the lipid uptake and accumulation in CD8^+^ TILs.

The role of CD36 in OxLDL uptake has been well established in the progression of atherosclerosis (Jay et al., 2015; Kita et al., 2000; Mitra et al., 2011; Navab et al., 2004; Pepino et al., 2014). CD36-mediated OxLDL import by macrophages induces lipid accumulation, particularly of cholesterol, and triggers inflammation and apoptosis in macrophages and endothelial cells, which manifests as fatty streaks on artery walls and are notable features of advanced atherosclerosis (Boullier et al., 2001). Interestingly, we found that CD8^+^ TILs also import OxLDL and provide the first evidence that this induces lipid peroxidation and suppresses CD8^+^ T cell effector functions in a CD36-dependent manner. LDL-bound PUFAs are prone to ROS-mediated lipid peroxidation (Yang et al., 2016). It has been shown that OxLDL or OxPLs cause oxidative stress, inhibit mitochondrial activity, and induce lipid peroxidation in primary hepatocytes (Sun et al., 2020). However, OxLDL per se is likely not the only source of oxidized lipids because as the E06 staining noted, OxPLs were highly abundant in the TME and dying tumor cell vesicles and apoptotic bodies could be other sources of CD36-mediated uptake of oxidized lipids. Phospholipid peroxidation can damage lipid membranes and also trigger ferroptosis, which is characterized by iron-dependent accumulation of lipid hydroperoxides (Stockwell et al., 2017). GPX4 is a known phospholipid hydroperoxidase that protects against lipid peroxidation-induced ferroptosis in tumor cells (Dixon et al., 2012; Yang et al., 2014), and plays a role in maintaining periphery homeostasis and antigen-stimulated proliferation of T cells (Matsushita et al., 2015). Not only did we show that OxLDL was co-localized with CD36, but we demonstrated that GPX4 over-expression restores CD8^+^ T cell function *in vivo*. This generates a model whereby known CD36 ligands found in the TME, like OxLDL and OxPLs, are internalized and trigger phospholipid peroxidation in CD36-expressing T cells, which can be ameliorated by enhanced GPX4 expression. A mechanistic gap in this model is how the increased lipid peroxidation suppresses CD8^+^ TIL effector functions because *Cd36^-/-^* CD8^+^ TILs had increased GZMB and cytokine production. We postulate that increased PLs peroxidation affects the plasma membrane and lipid rafts in a way that suppresses TCR signaling (Agmon et al., 2018; Zech et al., 2009).

Our study unveils a new mode of immunosuppression in the TME, opening previously unappreciated links to explore between lipid oxidation and cancer immunotherapy. The substantial increases in lipid peroxidation in CD8^+^ TILs could induce their ferroptosis locally in the TME, limiting their numbers, but these aspects of the model remain to be more deeply investigated. As our work suggests the role of GPX4 in regulation of anti-tumor functions of CD8^+^ TILs, therapeutic induction of ferroptosis in cancer cells needs to be approached with caution to avoid unwanted off-target effects on T cells. Recent studies show that genetic ablation or antibody-mediated blockade of CD36 retarded tumor growth (Al-Khami et al., 2017; Su et al., 2020; Wang et al., 2020). Our study presents the first evidence that CD36 acts in a CD8^+^ T cell autonomous manner to suppress TIL cytokine production and granzyme B expression, and genetic ablation of CD36 reduces melanoma growth in a T cell-dependent manner. Consistently, we also show that systemic blockade of CD36 with blocking mAb reduces tumor burden (Wang et al., 2020). Interestingly, CD36 is a marker for some types of cancer stem cells, and cancer-specific ablation of CD36 suppresses murine leukemia growth, and metastasis of murine glioblastoma or human oral cancer (Hale et al., 2014; Pascual et al., 2017; Ye et al., 2016). Taken together, these data suggest that CD36 is a promising immunotherapeutic target, acting on many cell types in a concerted manner to impair anti-tumor immune responses whilst simultaneously enhancing tumor progression and spread.

## STAR Methods

### KEY RESOURCES TABLE

| REAGENT RESOURCE | SOURCE | IDENTIFIER |
| --- | --- | --- |
| **Antibodies** |  |  |
| Brilliant Violet 711™ anti-mouse CD45 | BioLegend | Cat# 103147 |
| APC-eFluor® 780 anti-Mouse CD3 | Invitrogen | Cat# 47003282 |
| BUV737 anti-mouse CD4 | BD | Cat# 612761 |
| BUV395 anti-mouse CD8a | BD | Cat# 563786 |
| V500 anti-mouse CD44 | BD | Cat# 560781 |
| Brilliant Violet 605 anti-mouse PD-1 | BioLegend | Cat# 135220 |
| Brilliant Violet 711 anti-mouse Tim-3 | BioLegend | Cat# 119727 |
| Pacific Blue anti-mouse CD45.1 | BioLegend | Cat# 110722 |
| PerCP-Cy5.5 anti-mouse CD45.2 | BioLegend | Cat# 109828 |
| Brilliant Violet 421 anti-rat CD90/mouse CD90.1 | BioLegend | Cat# 202529 |
| PerCP/Cyanine5.5 anti-mouse CD90.2 | BioLegend | Cat# 105338 |
| PE anti-mouse CD36 | BD | Cat# 562702 |
| APC anti-mouse FOXP3 | Invitrogen | Cat# 1936927 |
| PE/Cy7 anti-mouse IFN-γ | BioLegend | Cat# 505826 |
| PE anti-mouse TNF | BioLegend | Cat# 506306 |
| FTIZ anti-mouse GZMB | BioLegend | Cat# 515403 |
| Anti-mouse CD16/32 | BioLegend | Cat# 101302 |
| PE anti-mouse TOX | Invitrogen | Cat# 12-6502-82 |
| PerCP-Cy5.5 anti-human CD3 | BioLegend | Cat# 300430 |
| Alexa Fluor® 647 anti-human CD4 Antibody | BioLegend | Cat# 344636 |
| Brilliant Violet 421™ anti-human CD45RA Antibody | BioLegend | Cat# 304130 |
| APC/Cyanine7 anti-human CD197 (CCR7) Antibody | BioLegend | Cat# 353212 |
| Brilliant Violet 605™ anti-human CD279 (PD-1) Antibody | BioLegend | Cat# 329924 |
| Brilliant Violet 421™ anti-human CD36 | BioLegend | Cat# 336230 |
| APC anti-human CD8a | Invitrogen | Cat# 17-0087-42 |
| PE-Cy™ 7 Mouse Anti-Human IFN-γ | BD | Cat# 560924 |
| APC anti-human TNF | Invitrogen | Cat# 17-7349-41 |
| **Buffers, chemicals, Peptides, and Recombinant Proteins** |  |  |
| penicillin-streptomycin | Invitrogen | Cat# 15140148 |
| G418 Sulfate | Invitrogen | Cat# 10131027 |
| DMEM | GIBCO | Cat# 11965118 |
| DMEM/F-12 | GIBCO | Cat# 11320082 |
| PBS | GIBCO | Cat# 20012-068 |
| RPMI 1640 Medium | GIBCO | Cat# 21875034 |
| Non-essential amino acids | GIBCO | Cat# 11140-035 |
| Fetal bovine serum | Sigma-Aldrich | Cat# F0804 |
| 2-Mercaptoethanol | GIBCO | Cat# 21985023 |
| L-Glutamine | GIBCO | Cat# 25030081 |
| Recombinant Murine IL-2 | PEPROTECH | Cat# 212-12 |
| DnaseI | Sigma | Cat# DN25-5G |
| 10× Permeabilization buffer | Invitrogen | Cat# 00-8333-56 |
| BD Cytofix/Cytoperm | BD | Cat# 554714 |
| eBioscience<br>Foxp3/Transcription factor<br>staining buffer set | eBioscience | Cat# 00-5523-00 |
| GP (33-41) peptide | GenScript | Cat# RP20091 |
| ACK buffer | Thermo | Cat# A1049201 |
| Percoll | GE Healthcare | Cat# 17-0891-01 |
| Colagenase | Sigma-Aldrich | Cat# 11088882001 |
| OxLDL | Kalenbiomed | Cat# 770252-7 |
| LDL | Kalenbiomed | Cat# 770200-4 |
| HDL | Kalenbiomed | Cat# 770300-4 |
| PMA<br>(Phorbol 12-myristate 13-acetate) | Sigma | Cat# P8139-1MG |
| Ionomycin | Sigma | Cat# 10634-1MG |
| Brefeldin A | BioLegend | Cat# 420601 |
| Other resources |  |  |
| 70 µM cell strainers | Corning Falcon | Cat# 352350 |
| Single Cell 3' Library &<br>Gel Bead Kit v2 | 10X Genomics | Cat# PN-120237 |
| Experimental Models:<br>Organisms/Strains |  |  |
| Mouse: C57BL/6J | Jackson | Strain# 027 |
| Mouse: *Cd36*<sup>-/-</sup> | (Coburn et al.,<br>2000) | JAX Strain# 019006 |
| Mouse:<br>Tg(TcrLCMV)327Sdz<br>(P14) | (Pircher et al.,<br>1989) | MGI:2665105 |
| Mouse: 10BiT | (Kaech et al.,<br>2003) | MGI:3767675 |
| Software and Algorithms |  |  |
| Flowjo | FlowJo, LLC | https://www.flowjo.com/ |
| R 3.6.1 | CRAN | https://cran.r-project.org/mirrors.html |
| RStudio 1.2 | RStudio | https://rstudio.com/ |
| IDEAS V6 | Aminis | https://www.luminexcorp.com/ |
| Seurat V3 | (Stuart et al.,<br>2019) | https://satijalab.org/seurat/v3.0/ |
| edgeR | (Robinson et al.,<br>2010) |  |
| Limma | (Ritchie et al.,<br>2015) |  |

### Mice

C57BL/6/J mice were purchased from Jackson Laboratories. Generation of *Cd36*^-/-^ mice were previously described (Coburn et al., 2000). 10BiT mice (Maynard et al., 2007) and P14 mice have been described (Kaech et al., 2003). P14 *Cd36*^-/-^ mice were generated by crossing P14 mice with *Cd36*^-/-^ mice. Animals were housed in specific-pathogen-free facilities at the Salk Institute and all experimental studies were approved and performed in accordance with guidelines and regulations implemented by the Salk Institute Animal Care and Use Committee.

### Cell lines and cell cultures

B16-F10 melanoma cell line (B16) was cultured in DMEM with 10% fetal bovine serum and 1% penicillin-streptomycin (Invitrogen). B16-F10 melanoma cell line that expresses gp33 (B16-gp33), was cultured in DMEM with 10% fetal bovine serum, 1% penicillin-streptomycin and 250 μg/ml G418 (Invitrogen #10131027). B16 and B16-gp33 were a gift from Hanspeter Pircher (University of Freiburg, Germany) (Prevost-Blondel et al., 1998). MC38 colon adenocarcinoma cell line was maintained in DMEM/F12 medium with 10% fetal bovine serum, 1% penicillin-streptomycin and MEM Non-Essential Amino Acids. All the tumor cell lines were used for implantation when in exponential growth phase.

For *in vitro* T cell culture, P14 splenocytes were activated in RPMI 1640 medium containing 10% fetal bovine serum and 1% penicillin-streptomycin, 2 mM L-glutamine, 0.1 μg/ml gp33 and 10 U/ml IL-2. At 24 hrs post activation,_P14 splenocytes were then treated with vehicle control (PBS), or 50 μg/ml OxLDL, or 50 μg/ml LDL, or 50 μg/ml HDL, or 100 μM SSO, or the combination of 100 μM SSO and 50 μg/ml OxLDL for another 48 hrs. To detect cytokine production *in vitro*, activated CD8 T cells were re-stimulated with 0.1 μg/ml gp33 in the presence 2.5 μg/ml Brefeldin A for 6 h at 37°C, followed by surface and intracellular staining described below.

For *ex vivo* CD8^+^ TILs cell culture, sorted CD8^+^ TILs were treated with vehicle control (PBS) or 50 μg/ml OxLDL for 24 hrs. To detect cytokine production *ex vivo*, CD8^+^ TILs were then stimulated with 50 ng/ml PMA and 3 μM Ionomycin in the presence 2.5 μg/ml Brefeldin A Solution for 4 h at 37°C, followed by surface and intracellular staining described below.

### Human PBMC *in vitro* culture

Buffy coats of healthy individuals were purchased from New York blood bank center. Peripheral blood mononuclear cells (PBMC) were isolated from buffy coats by density gradient cell separation. For this experiment, frozen PBMC from five different buffy coat were used. The frozen PBMC was washed and rested overnight for 16 hrs in RPMI 1640 medium with 10% fetal bovine serum and 1% penicillinstreptomycin at 37°C. After overnight rest, PBMC was washed and 10^5^ PBMC was plated per well. The PBMC was treated with either vehicle control (PBS), or OxLDL at a final concentration of 50 μg/mL (Invitrogen) for 3 days at 37°C and on day 4, SSO (Sigma-Aldrich) was added to the culture at a final concentration 100 μM. After 24 hours, the cells were stimulated with staphylococcal enterotoxin B (SEB) 5 μg/mL (List Biological Laboratories) and incubated for 16 hrs in the presence of Brefeldin A (Invitrogen), which was added 2 hours after the beginning of incubation.

### TIF and serum collection

TIF was collected from tumors using a previously described approach (Ho et al., 2015; Sullivan et al., 2019). Tumors were briefly rinsed in PBS and blotted on filter paper (VWR, Radnor, PA, 28298-020). The tumors were then put onto 70 μm cell strainers (VWR) affixed atop 50mL conical tubes, and centrifuged for 10 min at 4°C at 100 g. TIF was then collected from the conical tube, frozen in liquid nitrogen and stored at −80°C until further analysis. Blood was collected from the same animal via retro-orbital bleeding, and centrifuged at 845 g for 10 minutes at 4°C to separate serum, which was frozen in liquid nitrogen and stored at −80°C until further analysis.

### Fatty acid measurements using LC/MS

Lipids were extracted using a modified version of the Bligh-Dyer method (Bligh and Dyer, 1959). Briefly, TIF and serum samples were diluted in 1 mL PBS and shaken in a glass vial (VWR) with 1 mL methanol and 2 mL chloroform containing internal standards (^13^C_16_-palmitic acid, d7-Cholesterol) for 30s. The resulting mixture was vortexed for 15s and centrifuged at 2400 × g for 6 min to induce phase separation. The organic (bottom) layer was retrieved using a Pasteur pipette, dried under a gentle stream of nitrogen, and reconstituted in 2:1 chloroform:methanol for LC/MS analysis. Lipidomic analysis was performed on an Vanquish HPLC online with a Q-Exactive quadrupole-orbitrap mass spectrometer equipped with an electrospray ion source (Thermo). Data was acquired in positive and negative ionization modes. Solvent A consisted of 95:5 water:methanol, solvent B was 60:35:5 isopropanol:methanol:water. For positive mode, solvents A and B contained 5 mM ammonium formate with 0.1% formic acid; for negative mode, solvents contained 0.028% ammonium hydroxide. A Bio-Bond (Dikma) C4 column (5 μm, 4.6 mm × 50 mm) was used. The gradient was held at 0% B between 0 and 5 min, raised to 20% B at 5.1 min, increased linearly from 20% to 100% B between 5.1 and 55 min, held at 100% B between 55 min and 63 min, returned to 0% B at 63.1 min, and held at 0% B until 70 min. Flow rate was 0.1 mL/min from 0 to 5 min, 0.4 mL/min between 5.1 min and 55 min, and 0.5 mL/min between 55 min and 70 min. Spray voltage was 3.5 kV and 2.5 kV for positive and negative ionization modes, respectively. Sheath, auxiliary, and sweep gases were 53, 14 and 3, respectively. Capillary temperature was 275°C. Data was collected in full MS/dd-MS2 (top 5). Full MS was acquired from 100–1500 m/z with resolution of 70,000, AGC target of 1×10^6^ and a maximum injection time of 100 ms. MS2 was acquired with resolution of 17,500, a fixed first mass of 50 m/z, AGC target of 1×10^5^ and a maximum injection time of 200 ms. Stepped normalized collision energies were 20, 30 and 40%.

Lipid identification was performed with LipidSearch (Thermo). Mass accuracy, chromatography and peak integration of all LipidSearch-identified lipids were verified with Skyline (MacLean et al., 2010). Peak areas were used in data reporting, data was normalized using internal standards.

### Tumor engraftment and treatment of tumor-bearing mice

For tumor engraftment, 10^5^ B16, or 10^5^ MC38 tumor cells were injected subcutaneously in 100μl PBS. Tumors were measured every 2-3 days post tumor engraftment with or without the indicated treatments and tumor volume was calculated by volume = (length × width^2^)/2. For antibody-based treatment, tumor-bearing mice were i.p. treated with α-CD4 antibody (200 μg per injection, clone GK1.5, BioXcell) and α-CD8 antibody (200 μg per injection, clone TIB-210, BioXcell) every three days from day 7 post tumor implantation, or α-CD36 antibody (200 μg per injection, clone CRF D-2717 (Wang et al., 2020)) every two days from day 7 post tumor implantation.

### Tumor digestion and cell isolation

Tumors were minced into small pieces in RPMI containing 2% fetal bovine serum, 0.5 μg/ml DNase I (Sigma-Aldrich), and 0.5 mg/ml collagenase Type I (Sigma-Aldrich) and kept for digestion for 30 min at 37°C, followed by filtration with 70 μm cell strainers. Filtered cells were incubated with ACK lysis buffer (Invitrogen) to lyse red blood cells, mixed with excessive RPMI 1640 medium containing 10% fetal bovine serum and 1% penicillin-streptomycin, and centrifuged at 400g for 5 min to obtain singlecell suspension.

### Uptake of fatty acids, cholesterol, or lipoproteins, neutral lipid content assay, lipid peroxidation assay, and mitochondrion staining

For measuring uptake of fatty acids or cholesterol, cells were incubated in PBS containing 0.5 μg/ml C1-BODIPY^®^ 500/510 C12 (ThermoFisher, D3823), or PBS containing 0.1 μg/ml BODIPY™ FL C16 (ThermoFisher, D3821) for 20 min at 37°C. For measuring uptake of cholesterol, cells were incubated in PBS containing NBD Cholesterol (ThermoFisher, N1148) at final concentration of 10 μM for 15 min at 37°C. For measuring LDL uptake, cells were incubated in PBS containing 0.3% BSA and 20 μg/ml BODIPY™ FL LDL (ThermoFisher, L3483) for 30 min at 37°C. For measuring OxLDL uptake, cells were incubated in PBS containing OxLDL-DyLight™-488 (1:20 dilution, Oxidized LDL Uptake Assay Kit, Cayman Chemical, #601180) for 30 min at 37°C. After incubation, cells were washed with FACS buffer for surface staining. For neutral lipid content detection, after permeabilization and fixation, cells were stained using BODIPY^®^ 493/503 (ThermoFisher, D3922) at a final concentration of 250 ng/ml together with other intracellular proteins. For measuring lipid peroxidation, cells were incubated in PBS containing 2 μM BODIPY^®^ 581/591 C11 reagent (ThermoFisher, C10445) for 30 min at 37°C before live dead and surface staining. Intratumoral CD8^+^ T cells or splenic CD8^+^ T cells were sorted prior to C11 lipid peroxidation assay to avoid interference of tumor cells in the assay. For measuring mitochondrial membrane potential, cells were washed and incubated with PBS containing 10 nM MitoTracker^®^ Deep Red FM (ThermoFisher) for 15min. After staining, the cells were washed and resuspended in fresh MACS buffer (PBS containing 2% FBS) for surface marker staining as described above.

### Flow cytometry, cell sorting and antibodies

Single cell suspensions were incubated with Fc receptor-blocking anti-CD16/32 (BioLegend) on ice for 10 min before staining. Cell suspensions were first stained with LIVE/DEAD^®^ Fixable Violet or Red Dead Cell Stain Kit (ThermoFisher) for 5 min at room temperature. Surface proteins were then stained in FACS buffer (PBS with 2% FBS) for 30 min at 4°C. To detect cytokine production *ex-vivo*, cell suspensions were re-suspended in RPMI 1640 containing 10% FBS, stimulated by 50 ng/ml PMA and 3 μM Ionomycin in the presence 2.5 μg/ml Brefeldin A (BioLegend #420601) for 4 h at 37°C. Cells were processed for surface marker staining as described above. For intracellular cytokine staining, cells were fixed in BD Cytofix/Cytoperm (BD #554714) for 30 min at 4 °C, then washed with 1× Permeabilization buffer (Invitrogen #00-8333-56). For transcription factor staining, cells were fixed in Foxp3 / Transcription Factor Fixation/Permeabilization buffer (Invitrogen #00-5521-00) for 30 min at 4 °C, then washed with 1 × Permeabilization buffer. Cells were then stained with intraceulluar antibodies for 30 min at 4 °C. Samples were processed on LSR-II flow cytometer (BD Biosciences) and data were analyzed with FlowJo V10 (TreeStar). Cells were sorted either on FACSAria™ III sorter or Fusion sorter (BD Biosciences). The following antibodies against mouse proteins were used: anti-CD45 (30-F11), anti-CD3ε (17-A2), anti-CD4 (GK1.5), anti-CD8a (53-6.7), anti-CD44 (IM7), anti-PD-1 (29F.1A12), anti-TIM-3 (RMT3-23), anti-Ly5.1 (A20), anti-Ly5.2 (104), anti-Thy1.1 (OX-7), anti-Thy1.2 (30-H12), anti-CD36 (CRF D-2712), anti-IgA (mA-6E1), anti-FoxP3 (FJK-16S), anti-IFN-γ (XMG1.2), anti-TNF-α (MP6-XT22), anti-GZMB (GB11). These antibodies were purchased from Invitrogen, Biolegend or eBiosciences.

To detect cytokine production *ex-vivo* in human PBMC, cell suspensions were re-suspended in RPMI 1640 containing 10% FBS, stimulated with Staphylococcal enterotoxin B (SEB) and incubated for 16 hrs in the presence of Brefeldin A. The cells were washed with PBS and stained with LIVE/DEAD™ Fixable Red Dead Cell Stain Kit (ThermoFisher) for 5 minutes at room temperature. Cells were then stained with surface antibodies CD3, CD4, CD8, CD45RA, CCR7, PD-1 and CD36 for 30 minutes at 4°C and cells were washed with FACS buffer. For intracellular staining, the PBMC were treated with Intracellular Fixation & Permeabilization Buffer Set (Invitrogen# 88-8824-00) for 20 minutes at 4°C. The PBMC were then incubated with IFN-γ and TNF antibody for 1 hour at 4°C. The PBMCs were washed with permeabilization buffer and fixed in 1% paraformaldehyde. Data was acquired on BD Fortesa and analyzed with Flowjo. Antibodies for flow cytometry against human CD3 (Clone UCHT1), CD4 (Clone SK3), CD45RA (Clone HI100), CCR7 (Clone CO43H7), PD-1 (Clone EH12.2H7), CD36 (Clone 5-271) were purchased from BioLegend; CD8 (Clone SK1), IFN-γ (Clone B27), from BD Biosciences and TNF (Clone MAb11) from Invitrogen.

### Human patient assessment

All patients signed an approved informed consent before providing tissue samples. Patient samples were collected on a tissue-collection protocol approved by the MSK Institutional Review Board. Single cell suspensions from patients’ tumors were obtained by digesting tumor samples with type I collagenase (2 mg/mL), type V hyaluronidase (2 mg/mL) and type IV deoxyribonuclease I (200 U/mL) in serum-free RPMI 1640 using a GentleMACS Octo Dissociator (Miltenyi Biotec). Human samples were analyzed following safety regulation and stained with the following antibodies for FACS analysis: anti-CD45 (2D1), anti-CD3 (SK7), anti-CD4 (SK3), anti-CD8 (RPA-T8), anti-CD36 (TR9), anti-PD1 (MIH4), and anti-TOX (TXRX10). Samples were acquired using a Cytek Aurora flow cytometer and data analyzed with FlowJo 10.6 software.

### Adoptive T cell transfer and retroviral transduction

10^6^ naïve gp33-specific *Cd36*^+/+^ or *Cd36*^-/-^ P14 TCR transgenic CD8^+^ cells were transferred to tumor-bearing mice 7 days post tumor engraftment (retro-orbital). For GPX4 over-expression, 293T cells were transfected with Eco-helper and either MSCV control vector or vector over-expressing GPX4. 48 hr later, supernatant containing retroviral particles was ready for transduction. P14 donor splenocytes were *in vitro* activated by 0.1 μg/ml gp33 and 10 U/ml IL-2 at 37°C for 24h, then spin-transduced (1500 g) with fresh RV supernatant from 293T cells for 90 min at 30 °C in the presence of 5 μg/ml polybrene. Right after viral transduction 5×10^5^ P14 congenic CD8^+^ T cells were transferred into C57BL/6 mice that were implanted with B16-gp33 cells 7 days ago.

### Polychromatic imaging cytometry

For Imagestream (Amnis) analysis, CD8^+^ T cells stained with antibodies as above were sorted. Single stained cells were used as compensation controls. Images were captured at 60 × magnification and analysis was performed using IDEAS v6 software (Amnis).

### Histology and immunostaining of oxidized phospholipids

Tissues were fixed in formalin for 24 hrs, and then dehydrated in 70% EtOH for 24 hrs before further embedding in molten paraffin wax. Paraffin sections were completed at the Tissue Technology Shared Resource at the UCSD Cancer Center. Sections were blocked sequentially by donkey serum and biotin/avidin blocking, and incubated with biotinylated E06 for 12 hrs at 4°C, followed by alkaline phosphatase conjugated avidin for 30 min at RT. The nuclei were stained with hematoxylin. Stained tissue was visualized with NanoZoomer Slide Scanner.

### Single-cell RNA sequencing

Sorted cells were partitioned into an emulsion of nanoliter-sized droplets using a 10x Genomics Chromium Single Cell Controller and RNA sequencing libraries were constructed using the Chromium Single Cell 3’ Library & Gel Bead Kit v2 (10X Genomics, Cat# PN-120237). Briefly, droplets containing individual cells, reverse transcription reagents and a gel bead were loaded with poly(dT) primers that include a 16 base cell barcode and a 10 base unique molecular index (UMI). Reverse transcription reactions were engaged to generate barcoded full-length cDNA followed by the disruption of emulsions using the recovery agent and cDNA clean up with DynaBeads MyOne Silane Beads (Thermo Fisher Scientific, Cat# 37002D). Bulk cDNA was amplified, and indexed sequencing libraries were constructed using the reagents from the Chromium Single Cell 3’ v2 Reagent Kit. Libraries were sequenced on NextSeq Sequencing System (Illumina Cambridge).

### Single-cell RNA sequencing data processing and analysis

Cell Ranger (version 2.1.1) (from 10x genomics) was used to process Chromium single cell 3’ v2 RNA-seq output files. First, we generated fastq files for the Read1 for cell barcode and UMI and Read2 for transcript applying cellranger mkfastq (with default parameters). Second, we aligned the Read2 to the mouse genome (GRCm38/mm10) with cell ranger count (with default parameters). Further analysis was performed using Seurat package (version 3.1.5) in R (version 3.6.1) (Butler et al., 2018). Before performed analysis, we applied the following filtering step: only genes expressed in 3 or more cells were preserved and cells with less than 200 or more than 3000 unique expressed genes were discarded to exclude low-quality cells or cell doublets. Cells expressing more than 8% reads mapped to mitochondria genes were also discarded to exclude low-quality or dying cells. Gene expression matrix was normalized and natural log-transformed, and highly variable features were identified via standard Seurat workflow (Stuart et al., 2019). Then samples of *Cd36*^+/+^ and *Cd36*^-/-^ TILs were integrated to obtain an expression matrix comprising 14924 genes across 3400 cells (1561 *Cd36*^+/+^ TILs, and 1839 *Cd36^-/-^* TILs) for the rest of the analysis. Differentially expressed genes were identified using FindMarkers from Seurat package. Boxplots and Violin plots were performed using the combination of Seurat and ggplot.

The analyses of publicly available single-cell RNAseq data (GSE72056) (Tirosh et al., 2016) was performed in Seurat. Only genes expressed in 3 or more cells were preserved and cells with less than 200 or more than 7500 unique expressed genes were discarded. Gene expression matrix was normalized and natural log-transformed, and highly variable features were identified via standard Seurat workflow. Gene expression levels (E) in these data consist of log-normalized count via default Seurat algorithm. The analysis was restricted to PD-1-expressing CD8^+^ TILs based on the expression of *CD3D, CD8A* and *PDCD1* (E > 0) but not *CD4* (E =0). The cells showing no expression of *HAVCR2* or *CD36* (E=0) were also excluded in the analysis.

### Bulk RNAseq analysis

The transcriptome of naïve P14 CD8 T cells, P14 CD8^+^ TCF7^+^ TIM-3^-^ progenitor-like cells, CD8^+^ TCF7^-^ TIM-3^+^ terminally exhausted T cells was downloaded from NCBI GEO (GSE114631) (Siddiqui et al., 2019). The data was processed with the edgeR-Limma Package (Ritchie et al., 2015; Robinson et al., 2010).

### Fatty acid oxidation

Fatty acid oxidation (FAO) assay was performed as described previously (Ye et al., 2016). Briefly, equal numbers of sorted CD8^+^ T cells were plated in 96-well plates supplemented with FAO assay medium (RPMI 1640 medium containing 2% FBS, 10μM palmitic acid, 1% fatty acid free BSA (Sigma), 500μM carnitine (Sigma)). Cells were pulsed for 6 hours with 0.5μCi [9,10-^3^H(N)]-palmitic acid (Perkinelmer) and the medium was collected to analyze the released ^3^H_2_O, formed during cellular oxidation of [^3^H] palmitate. Briefly, medium was precipitated by 10% trichloroacetic acid (Sigma) and then supernatant was neutralized with 6N NaOH (Sigma) and loaded into ion exchange columns packed with DOWEX 1X2-400 resin (Sigma). The radioactive product was eluted with water and quantitated by liquid scintillation counting.

### Statistical analyses

Statistical analyses were performed using the two-tailed, unpaired, Student’s t-test unless otherwise specified. Each point represented a biological replicate and all data were presented as the mean ± SEM. The *P* values were represented as follows: \*\*\**P* < 0.001, \*\**P* < 0.01 and \**P* < 0.05.

## Author contributions

S.X., G.C., S.M.K. conceptualized, designed and supervised the research. S.X. performed experiments with assistance from P.R.M., Z.X., D.C., V.T., Y.F.. O.C. performed human PBMC experiments. R.Z. and T.M. performed human TILs analysis. X.S. and W.T. helped with immunostaining of oxidized phospholipid. A.F.M.P. performed lipidomics analysis. A.W., J.S.L, S.K.V., B.M. helped with gene expression analysis. M.D., R.M.E., N.A.A., J.D.W.. M.N.S., P.-C.H., J.L.W., and B.E. provided scientific input. S.X., S.M.K. prepared the manuscript.

## Declaration of interests

G.C. receives research funding from Bayer AG and Boehringer Ingelheim, but the funding is not relevant to the current study. J.L.W and X.S. are named inventors on patent applications or patents related to the use of oxidation-specific antibodies held by UCSD. All other authors declare no conflict of interest.

## Acknowledgement

We thank Dr. Marcus W. Bosenberg for discussion; Dr. Annelise G Snyder for graphical assistance; Dr. Anna-Maria Globig for assistance in RNAseq; Dr. Hubert Tseng for grant application; Drs. Thomas H Mann and Heather M McGee for manuscript review; C. O’Connor, Lara Boggeman and C. Fitzpatrick at the Salk FACS core; Nasun Hah and Tzu-Wen Wang at the Salk Sequencing core; the UCSD FACS core for cell sorting; Dr. M. Valeria Estrada at UCSD Histology core; and Faye McDonald for administrative assistance. We thank Dr. Yoav Altman from the Sanford Burnham Prebys Flow Cytometry Core, NCI grant P30 CA030199, and the James B. Pendleton Charitable Trust (helped purchase the Amnis) for Amnis analysis. The NGS Core Facility of the Salk Institute is supported with funding from NIH-NCI CCSG: P30 014195, the Chapman Foundation and the Helmsley Charitable Trust. The Razavi Newman Integrative Genomics and Bioinformatics Core is supported by NIH/NGMS R01 GM102491-07, NIH/NCI P30 CA014195-46, NIA/NMG 1RF1AG064049-01, and the Helmsley Trust. The Mass Spectrometry Core of the Salk Institute is supported with funding from NIH-NCI CCSG: P30 014195 and the Helmsley Center for Genomic Medicine. The MS data described here was gathered on a ThermoFisher Q Exactive Hybrid Quadrupole Orbitrap mass spectrometer funded by NIH grant (1S10OD021815-01). Tissue Technology Shared Resource is supported by a National Cancer Institute Cancer Center Support Grant (CCSG Grant P30CA23100). X.S. is supported by NIH grant K99HL148504. R.M.E is an investigator of the Howard Hughes Medical Institute and March of Dimes Chair in Molecular and Developmental Biology at the Salk Institute and supported by the NIH (DK057978, HL105278, ES010337), the Cancer Center (CA014195), a NOMIS Foundation Distinguished Scientist and Scholar Award, Lustgarten Foundation, the Don and Lorraine Freeberg Foundation. G.C. is supported by a Helmholtz Young Investigator Award (Helmholtz Gemeinschaft, #VH-NG-1113) and German Research Foundation (Deutsche Forschungsgemeinschaft, #CU375/7-1). This work was supported by NIH (R01CA240909 to S.M.K, R01 CA206483 to S.M.K and B.E., R01 CA206483 supplemental to P.R.M and R01HL148188 to X.S, and J.L.W.), and Melanoma Research Alliance (S.M.K, M.W.B), Genentech Foundation Fellowship (S.X.), and Salk Innovation Grant (S.M.K, R.M.E).

## Supplemental figure legends

**Figure S1.**
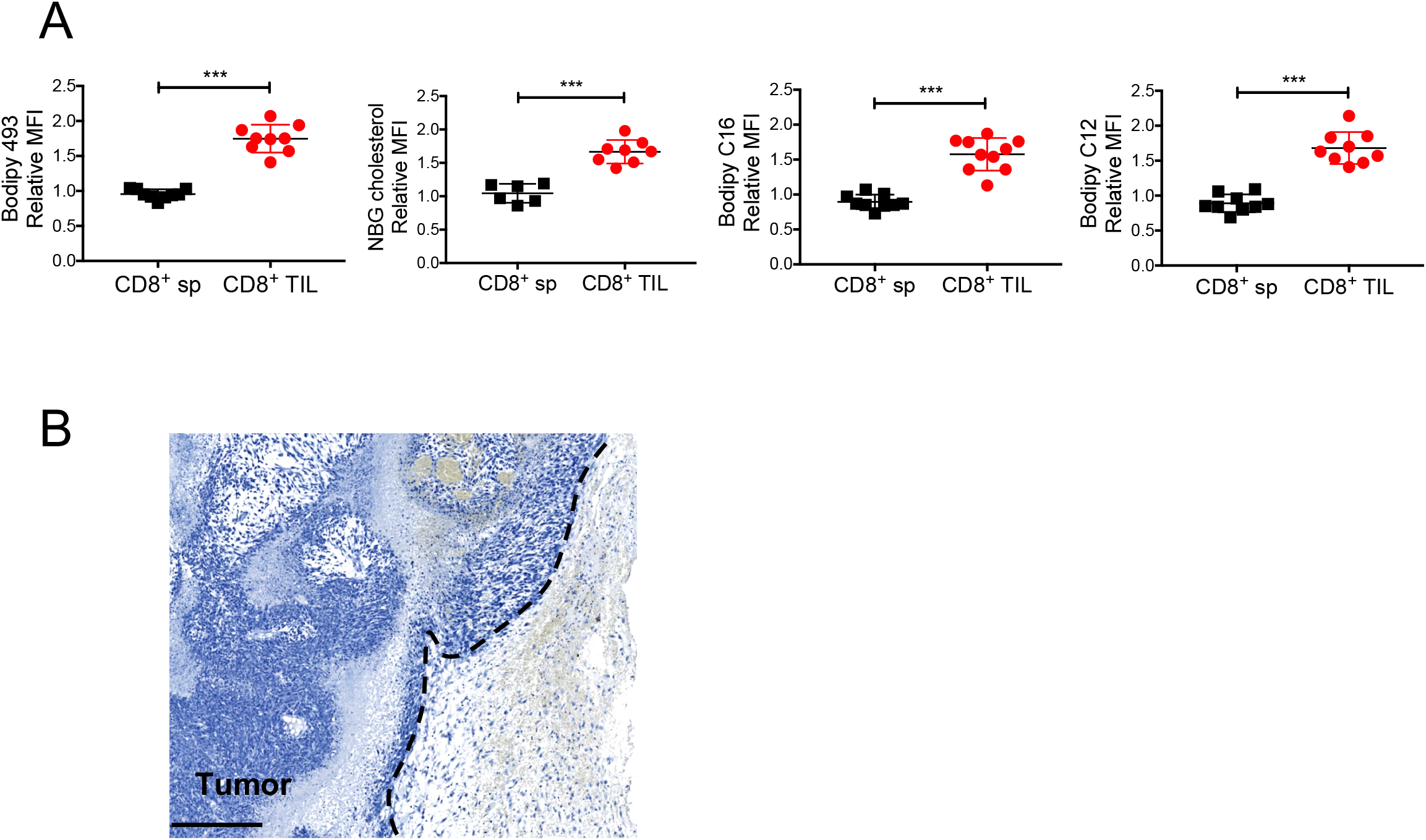
Increased lipid uptake and storage in CD8^+^ TILs in murine tumors. (A-B) C57BL/6J mice were implanted with MC38 (A) or B16 (B) tumor cells as indicated below and tumors or splenocytes were examined 21 days later. (A) Neutral lipid content (Bodipy 493), uptake of cholesterol and fatty acids (NBD cholesterol, Bodipy C12, and Bodipy C16) was compared between CD8^+^ T cells isolated from the spleen (CD8^+^ sp) or tumors (CD8^+^ TILs) by flow cytometry. Congenic Thy1.1 naïve splenocytes were spiked into each well to control for sample-to-sample variation and serve as an internal reference for normalizing Bodipy or NBD staining. Relative Mean Fluorescent Intensity (MFI) was calculated as the MFI ratio between the CD8^+^ T cells in the spleen or tumor relative to those in the internal reference. Data shown are mean ± SEM and statistical analysis was performed by two-tailed unpaired Student’s t-test, ***p < 0.001. All the results were pooled from 2-3 experiments with n=6-10. (B) Negative staining of B16 tumors by anti-mouse IgM secondary antibody in the absence of IgM E06 primary. The nuclei were stained with hematoxylin. Scale bar, 250 μm.

**Figure S2.**
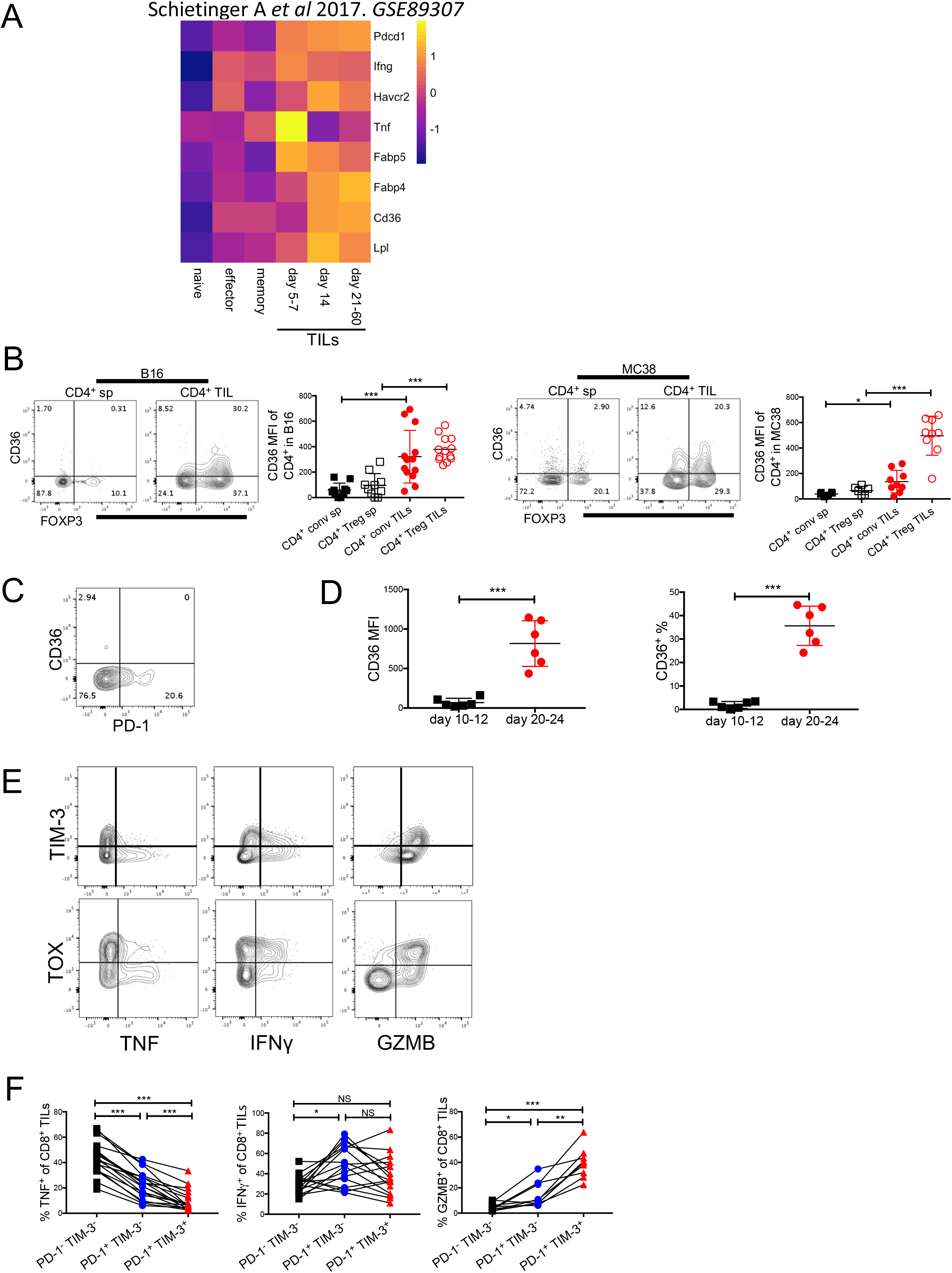
CD36 is expressed on functionally exhausted CD8^+^ TILs and regulatory CD4^+^ TILs. (B-F) C57BL/6J mice were implanted with B16 or MC38 tumor cells as indicated below, and tumors or splenocytes were examined 21 days later unless specified. (A) The transcript levels of indicated genes in naïve, effector, memory TCR-TAG CD8^+^ T cells (specific for SV40 large T antigen epitope I (TAG)) from TAG-expressing *Listeria monocytogenes*-infected spleens or tumor-reactive TCR-TAG CD8^+^ TILs isolated from liver cancer, were analyzed from the RNAseq dataset GSE89307 (Schietinger et al., 2016). The heatmap shows the mRNA expression normalized in row Z-score. (B) The expression of CD36 and FOXP3 was measured by flow cytometry in splenic CD4^+^ T cells (CD4^+^ sp) and CD4^+^ TILs from B16 tumors or MC38 tumors. (C) The expression of CD36 and PD-1 was measured by flow cytometry in CD8^+^ TILs from B16 tumors 12 days post implantation. (D) The expression of CD36 was measured by flow cytometry in CD8^+^ TILs from B16 tumors 10-12 days (early) vs. 20-24 days (late) post implantation. (E) The expression of TIM-3, TNF, IFNγ, GZMB, and TOX were measured in CD8^+^ TILs from B16 tumors. (F) The expression of TNF, IFNγ, or GZMB was measured in an effector cell subset (PD-1^-^ TIM-3^-^), an intermediate exhausted subset (PD-1^+^ TIM-3^-^), and a terminally exhausted subset (PD-1^+^ TIM-3^+^) of CD8^+^ TILs from B16 tumors. Data shown are mean ± SEM. Statistical analyses for (B, D, F) were performed by two-tailed unpaired Student’s t-test, NS, non-significant, *p < 0.05, **p < 0.01, ***p < 0.001. Samples were pooled from 2-3 experiments with each group containing n=6-9 (B), n=6 (D), n=9-18 (F) animals.

**Figure S3.**
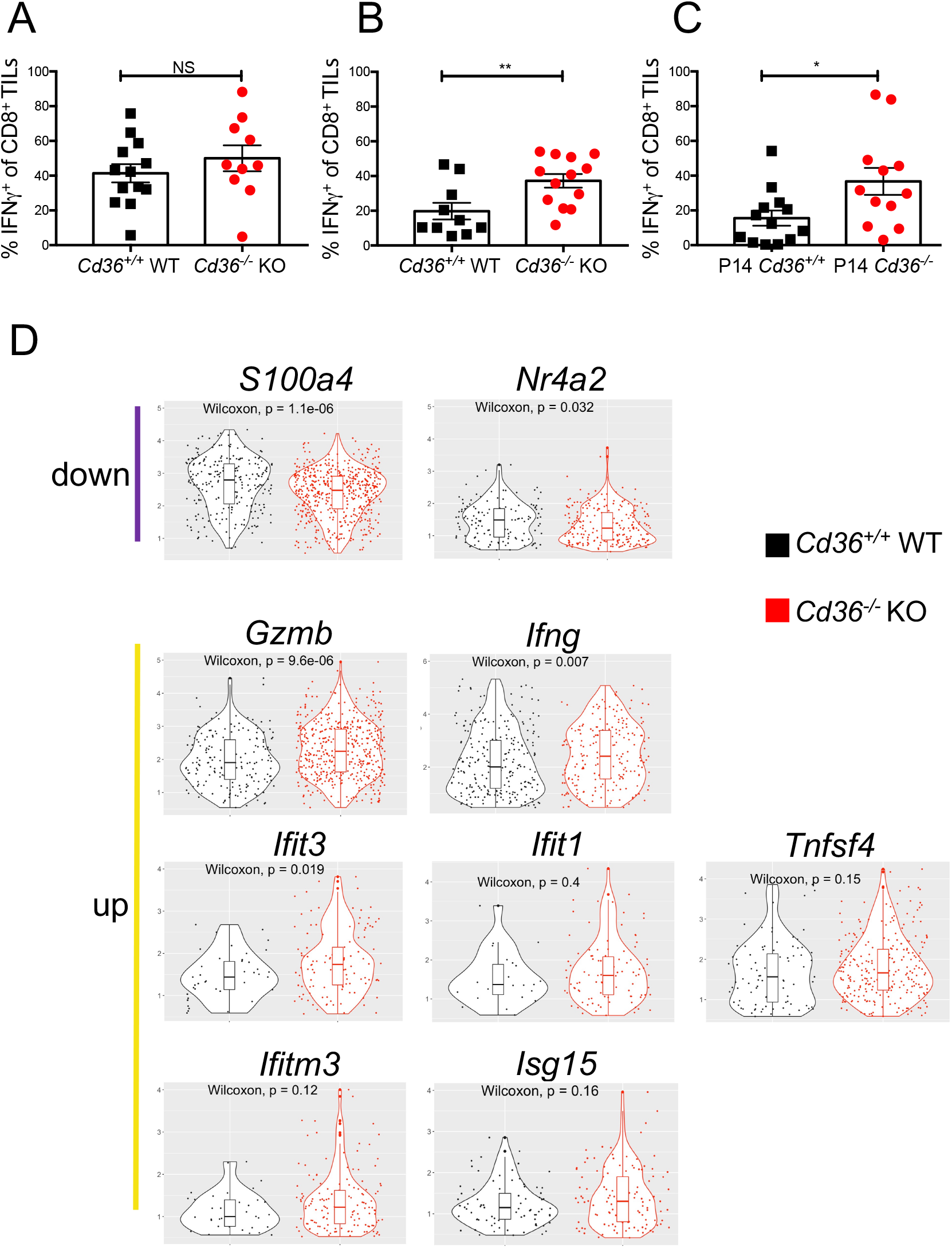
CD36 promotes CD8^+^ TIL dysfunction. (A-D) C57BL/6J mice were implanted with B16 or MC38 tumor cells as indicated below and tumors or splenocytes were examined 21 days later. (A-C) The expression of IFNγ was measured in *Cd36*^+/+^ or *Cd36^-/-^* CD8^+^ TILs from B16 tumors (A), MC38 tumors (B), P14 *Cd36*^+/+^ or *Cd36^-/-^* from B16 tumors (C). (D) scRNAseq was performed in *Cd36*^+/+^ or *Cd36^-/-^* TILs from B16 tumors. Violin plot + boxplot of differentially expressed genes were shown. The analyses in individual plots were restricted to cells whose expression of a given gene was great than 0. Data shown are mean ± SEM. Statistical analyses for (A-C) were performed by two-tailed unpaired Student’s t-test, NS, none-significant, *p < 0.05, **p < 0.01. Samples were pooled from 2-4 experiments with each group containing n=10-13 (A), n=10-13 (B), n=12-13 (C) animals.

**Figure S4.**
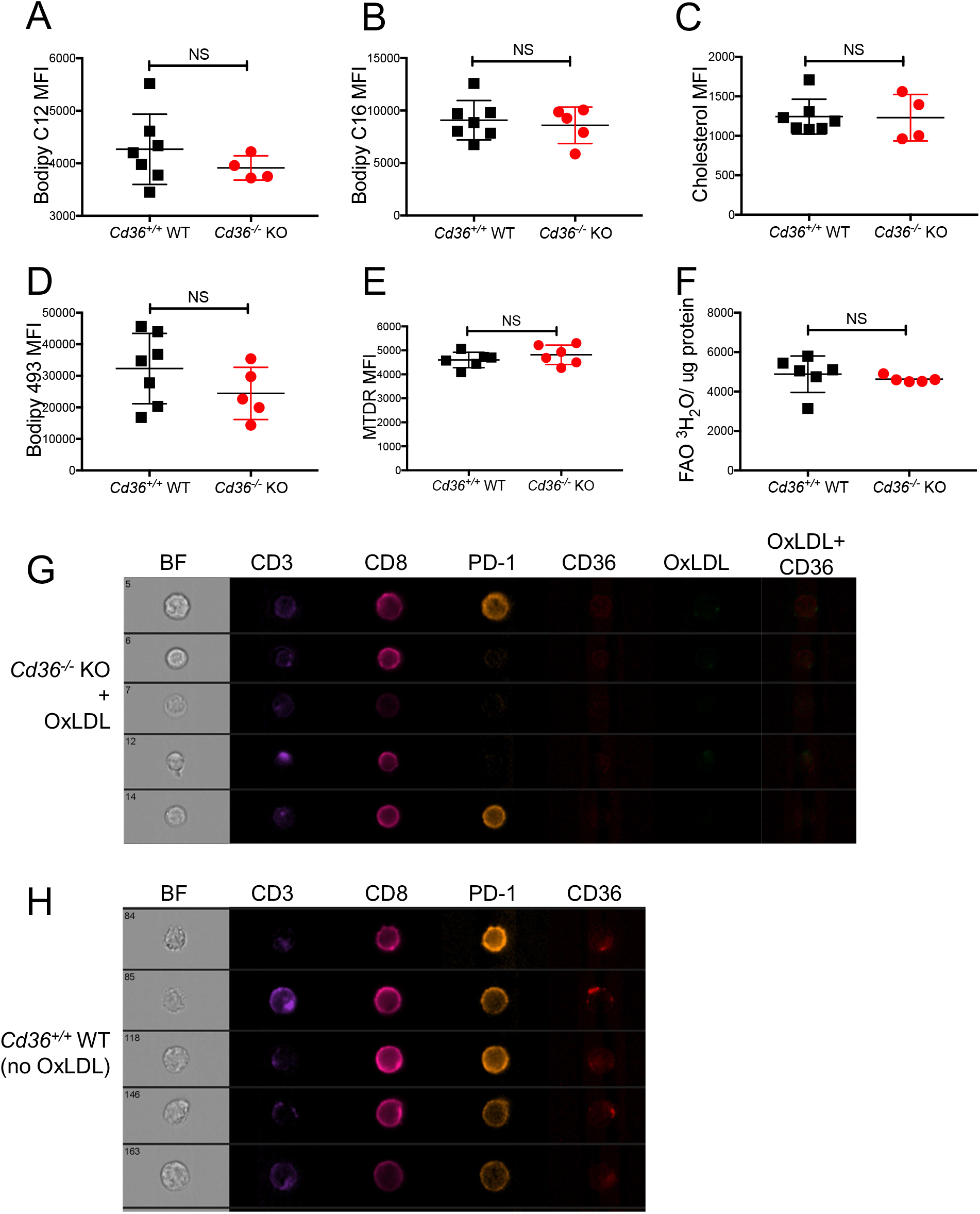
The impact of CD36 on metabolic activities of CD8^+^ TILs. (A-H) *Cd36*^+/+^ WT or *Cd36^-/-^* KO mice were were implanted with B16 tumor cells and tumors were examined 21 days later. (A-F) Uptake of Bodipy C12 (A), Bodipy C16 (B), uptake of cholesterol (C), neutral lipid content (D), mitochondrial potential (MitoTracker™ Deep Red FM) (E) and fatty acid oxidation (F) were measured in *Cd36*^+/+^ WT and *Cd36^-/-^* KO CD8^+^ TILs. (G-H) The expression of CD3, CD8, PD-1, CD36, and OxLDL uptake in *Cd36^-/-^* (G) and *Cd36*^+/+^ (H) CD8^+^ TILs were measured by Amnis ImageStream flow cytometry. Data shown are mean ± SEM. Statistical analyses for (A-F) were performed by two-tailed unpaired Student’s t-test, NS, none-significant. Samples were pooled from 2-3 experiments with each group containing n=4-7 (A), n=5-7 (B-C), n=4-6 (D), n=6 (E), and n=5-6 (F) animals.

**Figure S5.**
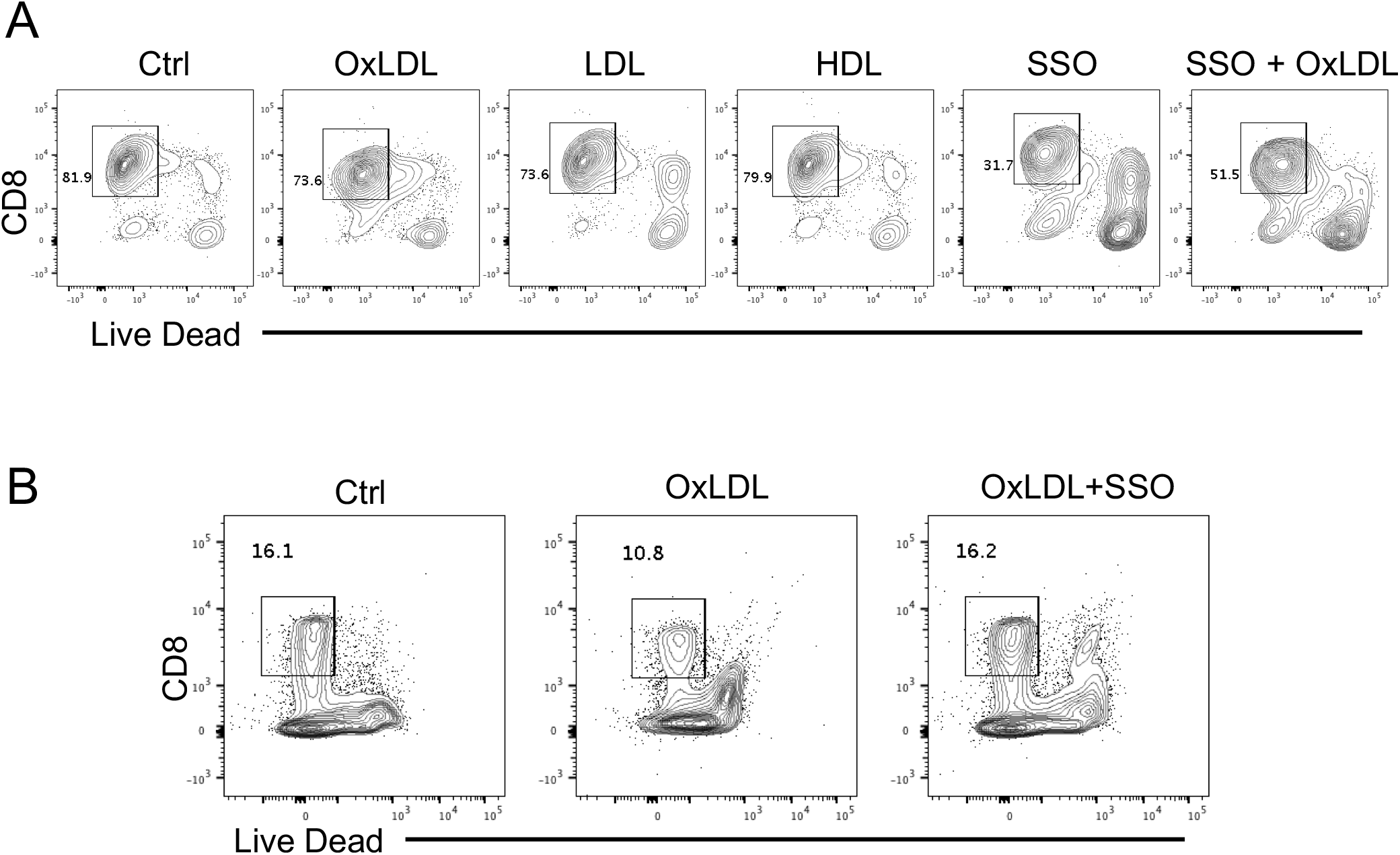
The impact of OxLDL on viability of CD8^+^ T cells *in vitro*. (A) P14 splenic CD8^+^ T cells were *in vitro* activated for 24 hrs, and then treated with vehicle control (Ctrl), OxLDL (50 μg/ml), LDL (50 μg/ml) or HDL (50 μg/ml), SSO (100 μM), or SSO + OxLDL for another 48 hrs. The cells were stimulated with gp33 peptide for 6 hrs. Viability was assessed via Live Dead assay by flow cytometry. (B) Human PBMCs were treated with either control (Ctrl), OxLDL (50 μg/ml), or the combination of oxLDL (50 μg/ml) and SSO (100 μM). Viability was assessed via Live Dead assay by flow cytometry.

